# Immunoproteasomal Processing of Isolevuglandin Adducts in Hypertension

**DOI:** 10.1101/2023.04.10.536054

**Authors:** Néstor de la Visitación, Wei Chen, Jaya Krishnan, Justin P. Van Beusecum, Venkataraman Amarnath, Elizabeth M. Hennen, Shilin Zhao, Mohammad Saleem, Mingfang Ao, David G. Harrison, David M. Patrick

## Abstract

Graphical Abstract

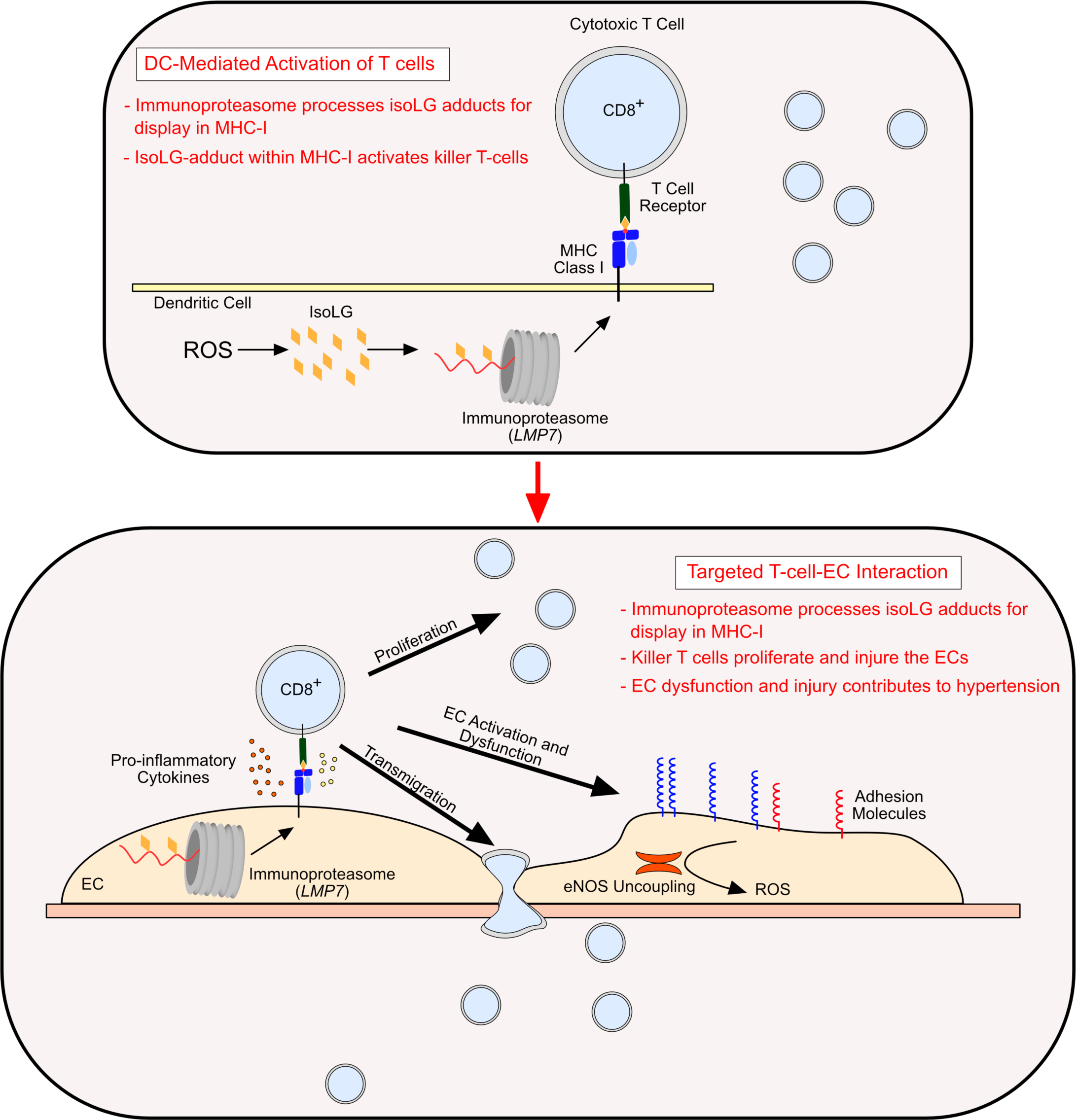

Isolevuglandins (isoLGs) are lipid aldehydes that form in the presence of reactive oxygen species (ROS) and drive immune activation. We found that isoLG-adducts are presented within the context of major histocompatibility complexes (MHC-I) by an immunoproteasome dependent mechanism.

Pharmacologic inhibition of LMP7, the chymotrypsin subunit of the immunoproteasome, attenuates hypertension and tissue inflammation in the angiotensin II (Ang II) model of hypertension. Genetic loss of function of all immunoproteasome subunits or conditional deletion of LMP7 in dendritic cell (DCs) or endothelial cells (ECs) attenuated hypertension, reduced aortic T cell infiltration, and reduced isoLG-adduct MHC-I interaction. Furthermore, isoLG adducts structurally resemble double-stranded DNA and contribute to the activation of STING in ECs. These studies define a critical role of the immunoproteasome in the processing and presentation of isoLG-adducts. Moreover they define a role of LMP7 as a regulator of T cell activation and tissue infiltration in hypertension.

## INTRODUCTION

Inflammation is a hallmark of hypertension.^1^ In response to hypertensive stimuli, antigen presenting cells, including macrophages and dendritic cells, gain the capacity to activate subsets of T cells.^2, 3^ Substantial evidence supports a role of CD8^+^ T cells in hypertension. In mice with Ang II induced hypertension, oligoclonal CD8^+^ T cells accumulate in the kidney and adoptive transfer of CD8^+^ T cells, but not CD4^+^ T cells restores hypertension in lymphocyte deficient *RAG-1^-/-^* mice. Youn et al. showed that even relatively young patients with hypertension exhibit an increase in circulating immunosenescent, proinflammatory, cytotoxic CD8^+^ T cells.^4^ Liu et al showed that high levels of CD8^+^ T cells accumulate in the kidney and stimulate activation of the sodium chloride co-transporter in mice with hypertension.^5^ Importantly, dendritic cells (DCs) from angiotensin II (Ang II) infused mice primarily support the proliferation and survival of CD8^+^ T cells.^6^

It is also likely that post-translationally modified proteins serve as antigens in hypertension. In particular, there is a striking increase in the formation of isolevuglandin (IsoLG)-adducted proteins within DCs of humans with hypertension and in experimental hypertension. IsoLGs are nonradical lipid aldehydes formed by enzymatic peroxidation of arachidonic acid and by free radical-mediated non-enzymatic peroxidation of prostaglandin H_2_ or H_2_-isoprostane. IsoLGs react rapidly with the primary amine of lysine residues on proteins thus forming isoLG adducts.^7^ DCs primed with isoLG-adducted proteins potently activate proliferation of CD8^+^ T cells, particularly if the latter are from hypertensive animals. Scavenging isoLGs with the small molecule 2-hydroxybenzylamine (2-HOBA) blunts hypertension in mice. DCs from hypertensive mice promote hypertension when adoptively transferred to naïve recipient mice, and this can be prevented if the donors are treated with 2-HOBA. Likewise, adoptive transfer of DCs in which isoLGs have been induced by exposure to tert-butyl hydroperoxide (tBHP) primes hypertension in naïve recipient mice.^6^

Priming of the immune system, however, does not fully explain the increase in vascular infiltration of immune cells observed in hypertension. Leukocyte and endothelial cell (EC) interactions are important drivers of immune activation and migration. ECs present antigens to CD8^+^ T cells in the context of major histocompatibility complexes (MHC-I). Lozanoska-Oscher *et* al. have shown that EC-specific MHC-I expression drives autoantigen specific CD8^+^ T cell transmigration.^8^ Pober *et* al. have shown that recognition of peptide antigens displayed in the context of MHC on the EC cell surface lead to T cell migration and activation.^9^ Importantly, ECs exhibit high levels of MHC-I expression highlighting their important role as directors of peripheral organ immune cell infiltration.^10^ The role of antigen presentation by ECs in the onset and development of hypertension is yet to be clarified.

CD8^+^ T cells are uniquely activated by antigens presented in the context of class 1 major histocompatibility complexes (MHC-I). The processing and packaging of such antigens initially requires proteasomal cleavage of proteins and their subsequent transfer to the endoplasmic reticulum, where they are loaded into MHC-I.^1^^1^ Once in the endoplasmic reticulum, peptides are further trimmed by endosomal proteases, and ultimately packaged into MHC-I that is then transported to the cell surface.^12^ Both self-peptides and peptides derived from foreign pathogens are displayed in this fashion for recognition by T cells and innate lymphocytes. Proteasomes exist as different subtypes, including the standard proteasome, the immunoproteasome, the thymoproteasome and the intermediate proteasome. These are characterized by differences in subunit enzymatic activity. The standard proteasome consists of three subunits, β1, β2, and β5.^13^ The 20S proteasome is primarily responsible for the generation of intracellular antigenic peptides that contribute to the MHC-I peptide pool.^12^ Inflammatory activation, primarily by interferons, induces the expression of the inducible subunits LMP2, MECL1, and LMP7 which combine to form the immunoproteasome.^14^ The different proteasome isoforms possess unique peptidase activities with the capacity to generate unique antigens.^15^

The above considerations are important for cells with high levels of oxidative stress in which isoLG formation occurs. High concentrations of isoLGs inhibit the proteolytic activity of standard proteasomes and even at very low concentrations isoLGs specifically inhibit the chymotrypsin-like activity of the 20S proteasome.^16^ Thus, the formation of isoLGs might paradoxically impair antigen presentation. Indeed, we have observed that presentation of the ovalbumin peptide SINFEKKL by DCs is impaired in mice with Ang II-induced hypertension, and this is restored to normal by scavenging isoLGs with 2-HOBA. Such observations are puzzling because the non-selective proteasome inhibitor Bortezomib (BTZ) attenuates Ang II induced hypertension and aortic remodeling in rats^17^ and reduces aortic aneurysm formation in ApoE mutant mice treated with this octapeptide.^18^ Importantly, mice mutant for LMP7, the chymotrypsin subunit of the immunoproteasome, are protected from Ang II-induced atrial fibrillation and cardiac hypertrophy.^19, 20^

In this study, we describe the intermolecular interaction of isoLG adducts with MHC-I in mouse DCs and ECs and demonstrate that this interaction is proteasome dependent. In Ang II treated mice, surface isoLG and isoLG-MHC-I interaction is restricted to CD11c^+^ in secondary lymphoid tissue and is uniquely present on ECs isolated from aorta. DCs from Ang II treated mice exhibit augmented proteasome activity and treatment of mice with BTZ attenuates Ang II-mediated hypertension.

Furthermore, we demonstrate an essential role of the immunoproteasome in the interaction of MHC-I with isoLG and the establishment of tissue inflammation and hypertension. Specifically, we describe a critical role of the chymotrypsin subunit LMP7 in Ang II mediated immune activation and hypertension in both DCs and ECs. These results implicate an essential role of the immunoproteasome in isoLG-mediated immune activation and hypertension and confirm a role of isoLGs as neoantigens presented within MHC-I.

## RESULTS

### IsoLG Adduct Interaction with MHC-I is Proteasome Dependent

In initial experiments we isolated splenic DCs from *C57Bl/6* mice and exposed these cells to the oxidant tBHP, which potently induces isoLG formation in these cells.^6^ In cells not exposed to tBHP, there was minimal surface isoLG-as evidenced by extracellular staining with the antibody D11, which detects isoLGs adducts independent of the peptide backbone. In contrast, exposure to this tBHP markedly increased surface isoLG-adducts. Next, we co-treated DCs with the proteasome inhibitors MG132 or BTZ and analyzed the intermolecular interaction of the MHC-I molecule H-2D^b^ with isoLG using fluorescence resonance energy transfer (FRET) by flow cytometry (Figure 1A). Importantly, all proteasome inhibitors abrogated both surface levels of isoLG-adducts and the association of these with H-2D^b^ in response to tBHP exposure (Figure 1B-1E). These data demonstrate that isoLG adducts accumulate on the surface of DCs, predominantly associated MHC-I, following an oxidant stimulus, in a proteasome-dependent fashion.

**Figure 1:**
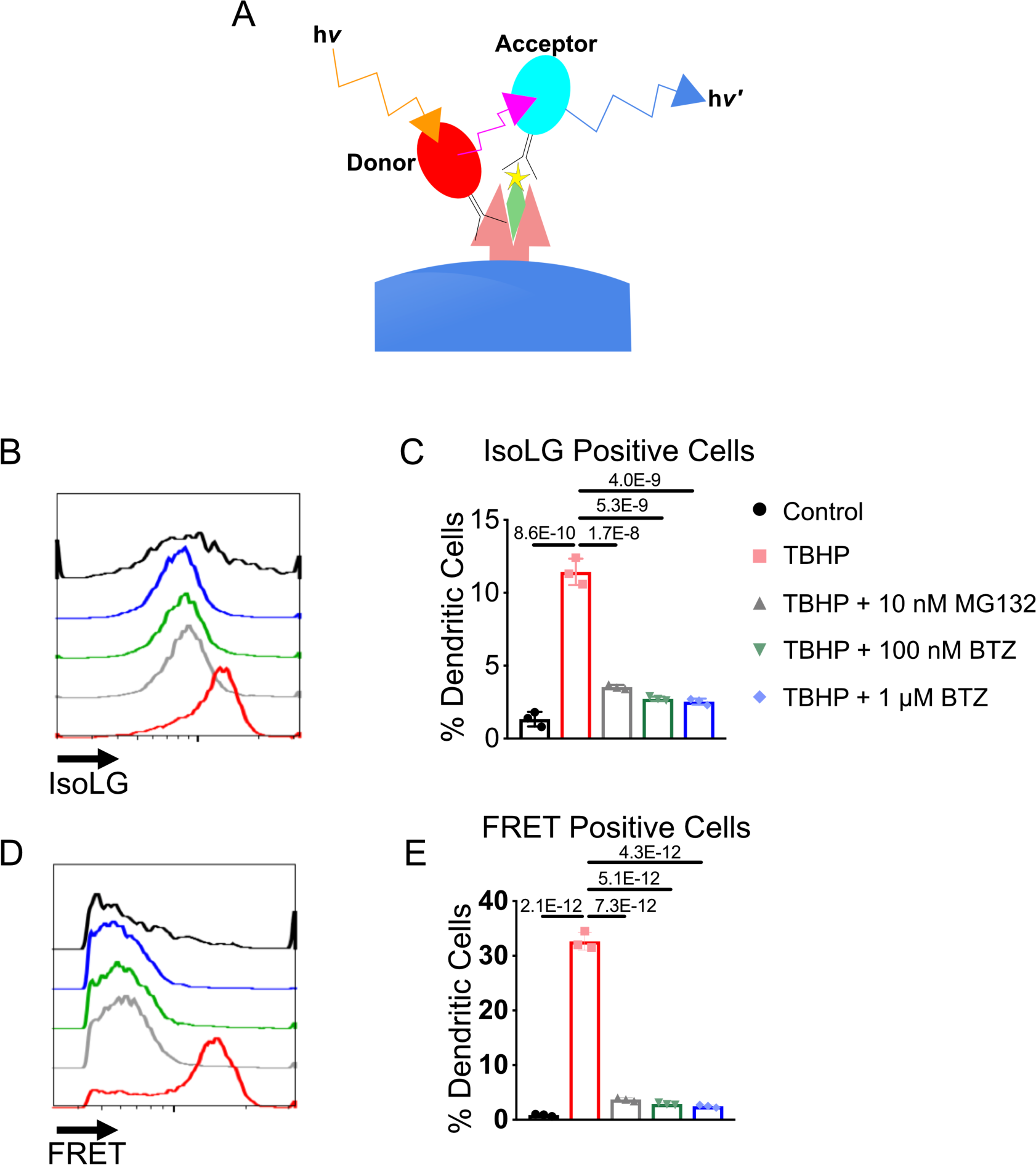
IsoLG-adduct interaction with H-2D^b^ is Dependent on Proteasome Activity. Intermolecular interaction of IsoLG-adducts with the MHC-Class I molecule H-2D^b^ was determined by FRET. **(A)** Paradigm of detection of isoLG-adduct H-2D^b^ interaction by FRET. A donor fluorophore is conjugated to an H-2D^b^ antibody and an acceptor fluorophore is conjugated to the D11 ScFv anti- isoLG adduct antibody. The donor fluorophore is excited by a laser at energy *hv* and the emission of the donor excites the acceptor fluorophore which emits photons at energy *hv’* that is detected by flow cytometry. **(B)** Representative histograms of surface isoLG-adducts from DCs treated with tBHP and proteasome inhibitors MG132 and BTZ. **(C)** Quantitation of isoLG positive DCs. **(D)** Representative histograms of FRET from DCs treated with tBHP. **(E)** Quantitation of FRET^+^ DCs. Data are representative of 3 repeated experiments. Data were analyzed by 1-way ANOVA with Tukey’s post-hoc test (*n* = 3).

### Hypertension induces Surface IsoLG Adduct Accumulation and Interaction with MHC-I in CD11c^+^ Antigen Presenting Cells

We have previously shown that there is a marked increase in isoLG-adduct formation in antigen presenting cells both experimental and human hypertension and a concomitant increase in superoxide production in DCs in experimental hypertension.^6^ Given our findings with oxidant exposure of DCs in culture, we hypothesized that hypertension *in vivo* would be associated with presentation of isoLG adducts in MHC-I. Thus, we infused *C57Bl/6* mice with 490 ng/kg/min of Ang II for 14-days and performed flow cytometry for surface isoLG-adducts and isoLG-adduct H-2D^b^ FRET. Using the gating strategy presented in Supplemental Figure 1A we found that Ang II-induced hypertension is associated with surface accumulation IsoLG-adducts and this is paralleled by an increase in association of these adducts with H-2D^b^ (Supplemental Figure 1B-E). Notably, these increases in MHC-I presentation of isoLG adducts was entirely in CD11c^+^ cells, confirming that surface isoLG is present exclusively on professional antigen presenting cells. The presentation of isoLG-adducts within MHC-I follows this pattern.

### Proteasome Activity and LMP7 Expression are Increased in APCs from Hypertensive Mice and Humans

The processing and ultimate presentation of antigens within MHC-I is critically dependent on proteasomal activity. In keeping with this, we found a significant increase in chymotrypsin-like activity in CD11c^+^ cells from spleens of Ang II-infused mice as compared to sham treated animals (Supplemental Figure 2A). To determine if proteasome activity contributes to Ang II-induced hypertension, we co-treated sham or Ang II infused mice with vehicle or 0.5 mg/kg of the non-selective proteasome inhibitor BTZ three times per week. While BTZ infusion did not affect blood pressure (BP) in sham infused mice, it reduced systolic blood pressure (SBP) in Ang II infused animals (Supplemental Figure 2B). Chymotrypsin-like proteasome activity was reduced in total splenocytes from BTZ treated animals confirming an efficient reduction in proteasome activity (Supplemental Figure 2C). The activities presented in Supplemental Figure 2A and 2C could reflect increases in either the standard proteasome or the immunoproteasome. A critical subunit of the immunoproteasome is LMP7 and immunoblot analysis of proteins isolated from DCs revealed Ang II-induced hypertension is associated with a striking increase this subunit. To determine if similar changes in LMP7 occur in human hypertension we analyzed proteins from CD14^+^ monocytes of normotensive or hypertensive subjects by immunoblot and found an almost 2-fold increase in LMP7 in hypertensive human monocytes (Supplemental Figure 2D-F, Supplemental Table 1). These findings confirm that DC specific proteasome activity is increased and contributes to Ang II-induced hypertension in mice. Furthermore, these results suggest an increase in immunoproteasome activity in murine and human hypertension, as reflected by an increased in LMP7 expression.

### LMP7 Inhibition Reduces Blood Pressure and Aortic Inflammation

We co-treated mice with Ang II infusion and vehicle or 6 mg/kg of the LMP7 specific inhibitor PR-957 three times weekly. Systolic blood pressure was reduced by PR-957 treatment (Figure 2A). Next, we performed isoLG-adduct-H-2D^b^ FRET from single cell suspensions prepared from the aorta. Importantly, we found a reduction in the association of MHC-I and isoLG-adducts in DCs from aortas harvested from PR-957 treated mice (Figure 2B and C). PR-957 treatment also reduced accumulation of CD3^+^ T cells (Figure 2D and E). Combined these results define an important role of LMP7 in Ang II-induced hypertension. Moreover, they confirm isoLG-adduct-MHC-I interaction in DCs at the site of inflammation. Finally, they define a role of LMP7 in T cell infiltration of the aorta in hypertension.

**Figure 2:**
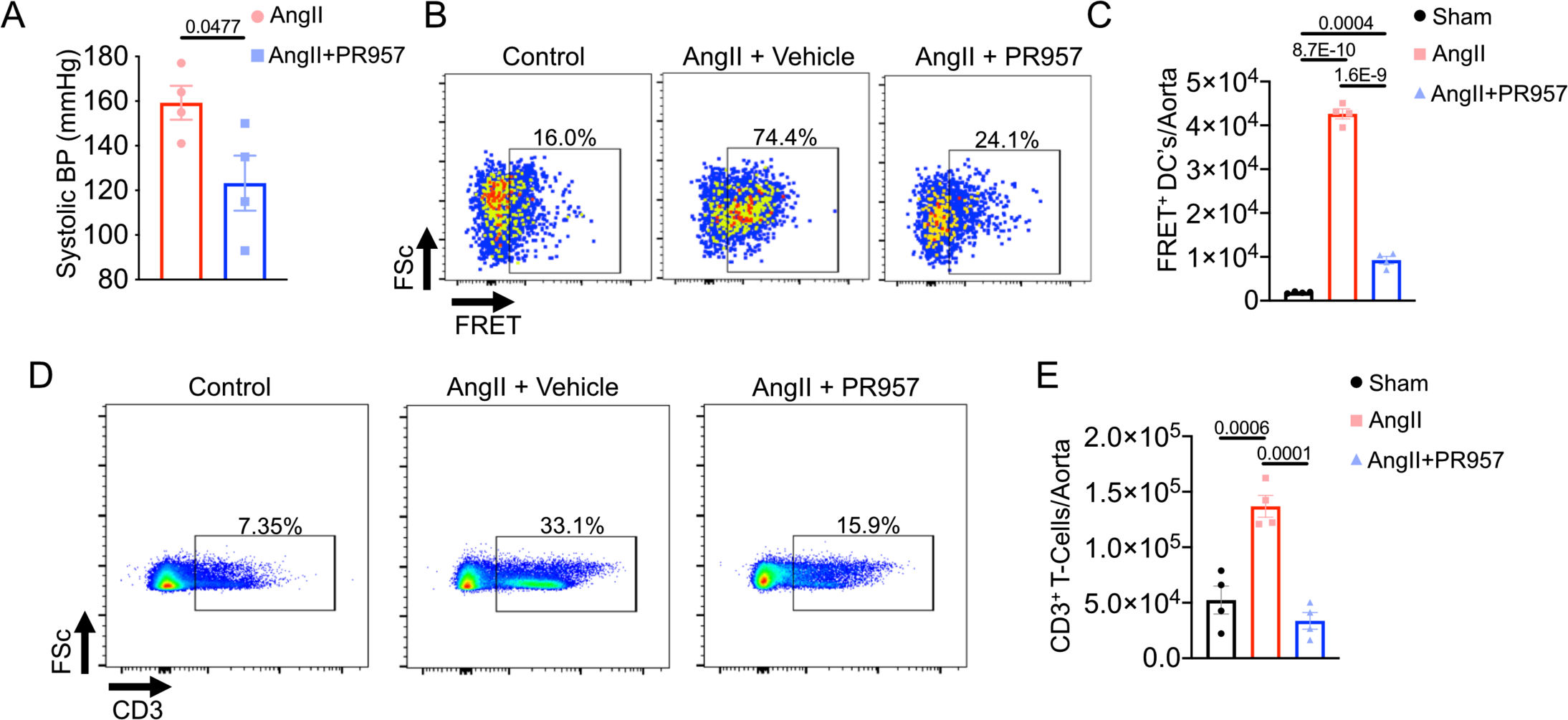
Inhibition of LMP7 attenuates hypertension and aortic inflammation. Mice were co-treated with angiotensin II and vehicle or PR-957 for 2-weeks. **(A)** Systolic blood pressure was measured. Data were analyzed by two-tailed Student’s T-test (*n* = 4). **(B)** Representative flow cytometry plots of isoLG-adduct-H-2D^b^ FRET in dendritic cells harvested from aorta. **(C)** Quantitation of isoLG-adduct-H-2D^b^ FRET^+^ dendritic cells harvested from aorta. **(D)** Representative flow cytometry plots of CD3^+^ T cells harvested from aorta. **(E)** Quantitation of CD3^+^ T cells harvested from aorta. Data were analyzed by 1-way ANOVA with Tukey’s post-hoc test (*n* = 4).

### Immunoproteasome Expression Increases IsoLG adduct-MHC-I Interaction

To gain further insight into the role of the immunoproteasome in processing isoLG adducts, we studied B8 cells, a mouse fibroblast line that expresses the standard proteasome but not immunoproteasome subunits.^21^ In addition, we also studied B8.27M.2 cells, which are stably transfected B8 cells that overexpress the immunoproteasome subunits LMP2, LMP7, and MECL2. We exposed B8 and B8.27M.2 cells with tBHP and performed isoLG-adduct-H-2D^b^ FRET for isoLG adduct-MHC-I interaction. At baseline, B8.27M.2 cells exhibited augmented isoLG adduct/MHC-I interaction as compared to the control B8 cells. Following tBHP treatment B8.27M.2 cells exhibited increased isoLG-adduct-H-2D^b^ FRET (Figure 3A-D), while B8 cells did not. These data support previous studies describing isoLG-mediated inhibition of standard proteasome activity. Moreover, they show that the immunoproteasome is not inhibited by isoLGs and that it is capable of processing isoLG adducted antigens that ultimately are loaded into MHC-I.

**Figure 3:**
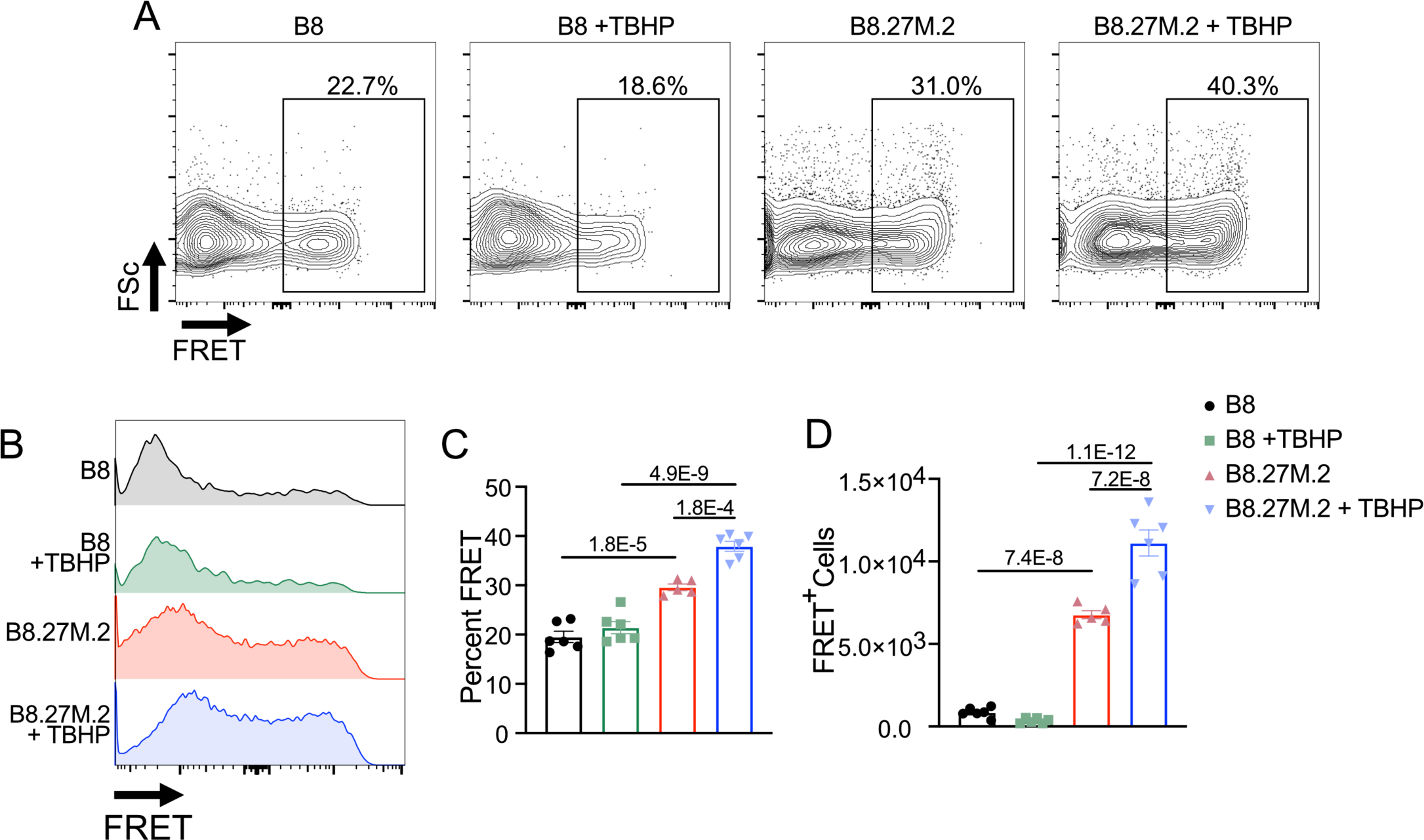
Overexpression of immunoproteasome subunits increases FRET. B8 and B8.27M.2 cells were co-treated with vehicle or tBHP. **(A)** Representative flow cytometry plots of FRET. **(B)** Representative histograms of FRET. **(C)** Quantitation of percent of FRET^+^ cells as a percentage of H2-D^b+^/IsoLG-adduct^+^ cells. **(D)** Quantitation of FRET^+^ cells. Data were analyzed using 1-way ANOVA with Tukey’s post-hoc test (*n* = 5-6).

### Loss of function of LMP7, LMP2, and MECL1 Attenuates Hypertension and Inflammation

In additional experiments, we studied *C57Bl/6* mice or mice lacking LMP7, LMP2, and MECL1 (triple knockout mice, TKO). Mice were infused with Ang II and blood pressure was measured by radiotelemetry. TKO mice are on the *C57Bl/6* background.^22^ The TKO mice exhibit an attenuated hypertensive response (Figure 4A-C). Moreover, TKO mice exhibit a reduction in renal CD8^+^ T cell infiltration compared to *C57Bl/6* wild-type mice (Figure 4D-F). Importantly, total renal T cells were reduced in the TKO mouse (Figure 4G). Finally, isoLG-adduct-H-2D^b^ FRET in DCs isolated from the aorta was significantly reduced in TKO mice (Figure 4H-J).

**Figure 4:**
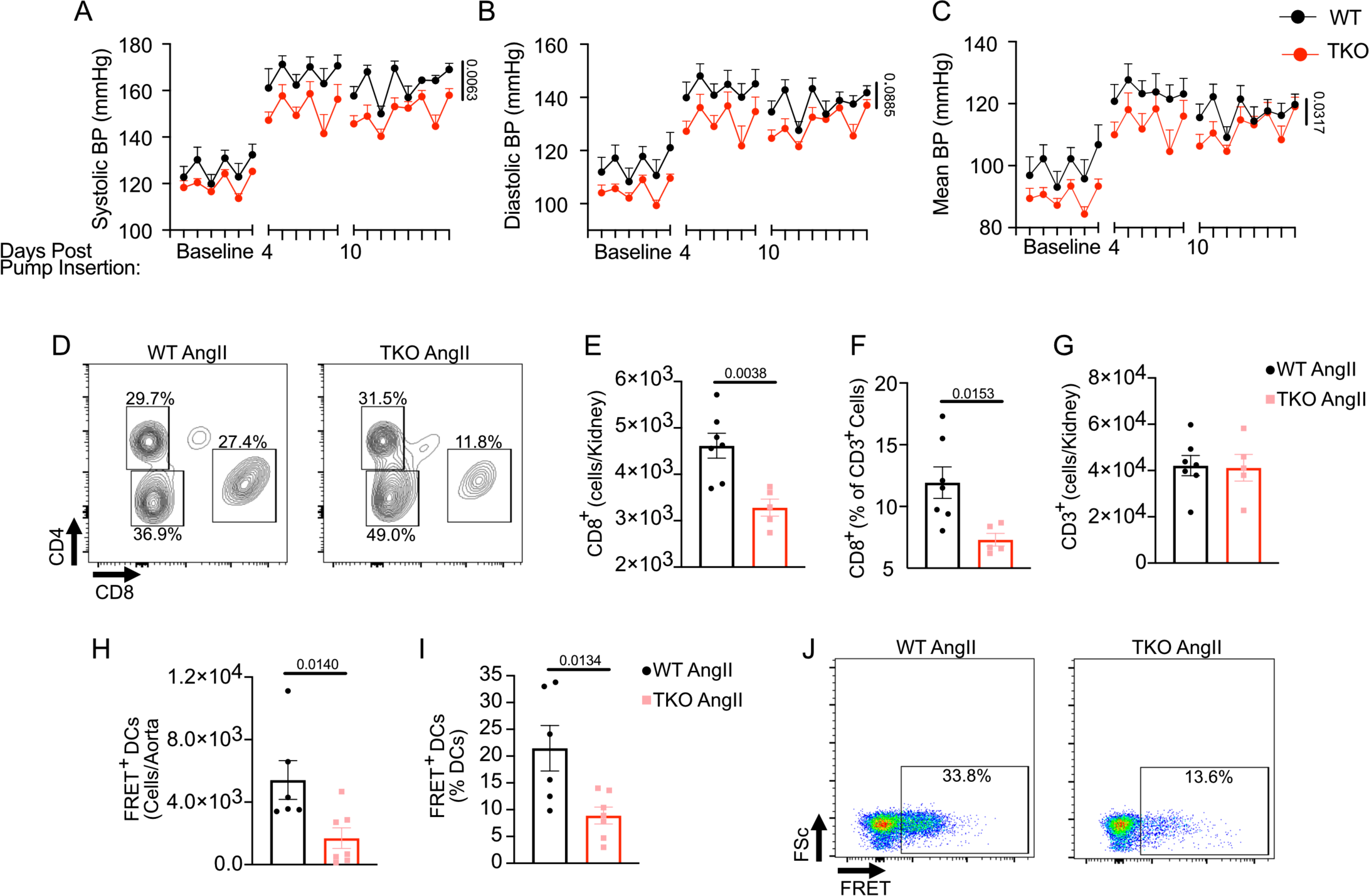
The immunoproteasome contributes to the development of hypertension. Immunoproteasome triple knockout mice (TKO) and control C57BL/6 mice were treated with angiotensin II. Radiotelemetry was used to measure **(A)** systolic, **(B)** diastolic, and **(C)** mean blood pressure. Blood pressure was analyzed using 2-way ANOVA (*n*=5-7). **(D)** Representative flow cytometry diagrams for CD4^+^ and CD8^+^ T cells harvested from Kidney. **(E)** Quantitation of CD8^+^ renal T cells. **(F)** Representation of CD8^+^ T cells as a percentage of total CD3^+^ T cells. **(G)** Quantitation of total CD3^+^ renal T cells. **(H)** Quantitation of FRET^+^ DCs harvested from aorta. **(I)** Representation of FRET^+^ aortic DCs as a percentage of total DCs. **(J)** Representative flow cytometry plots for FRET in DCs harvested from aorta. Data were analyzed by two-tailed Student’s T-test or Mann Whitney U Test (*n* = 6-7).

### Loss of Function of LMP7 in DCs Attenuates Hypertension and Inflammation

Using CRISPR/Cas9 we created mice with LoxP sites flanking exons 1 and 2 of the *Lmp7* locus in unconserved regions (Figure 5A). Targeting did not disrupt LMP7 expression in spleen, bone marrow, and kidney as measured by immunoblot (Figure 5B). Real-time RT-PCR performed with probes that overlap exons 1-2 and 4-5 from cDNA prepared from flow sorted CD11c^+^/IAb^+^ splenic DCs confirmed almost complete elimination of *Lmp7* mRNA in *Lmp7^fl/fl^/CD11c-Cre* DCs compared to *Lmp7^fl/fl^* DCs (Figure 5C and D). *Lmp7* and *Lmp7^fl/fl^/CD11c-Cre* mice were infused with Ang II and blood pressure was measured by radiotelemetry. *Lmp7^fl/fl^/CD11c-Cre* mice exhibited attenuated hypertension (Figure 5E and F; Supplemental Figure 4). Moreover, *Lmp7^fl/fl^/CD11c-Cre* mice exhibited a reduction in aortic DC isoLG-adduct-H-2D^b^ FRET and aortic T-cell infiltration (Figures 5G-I).

**Figure 5:**
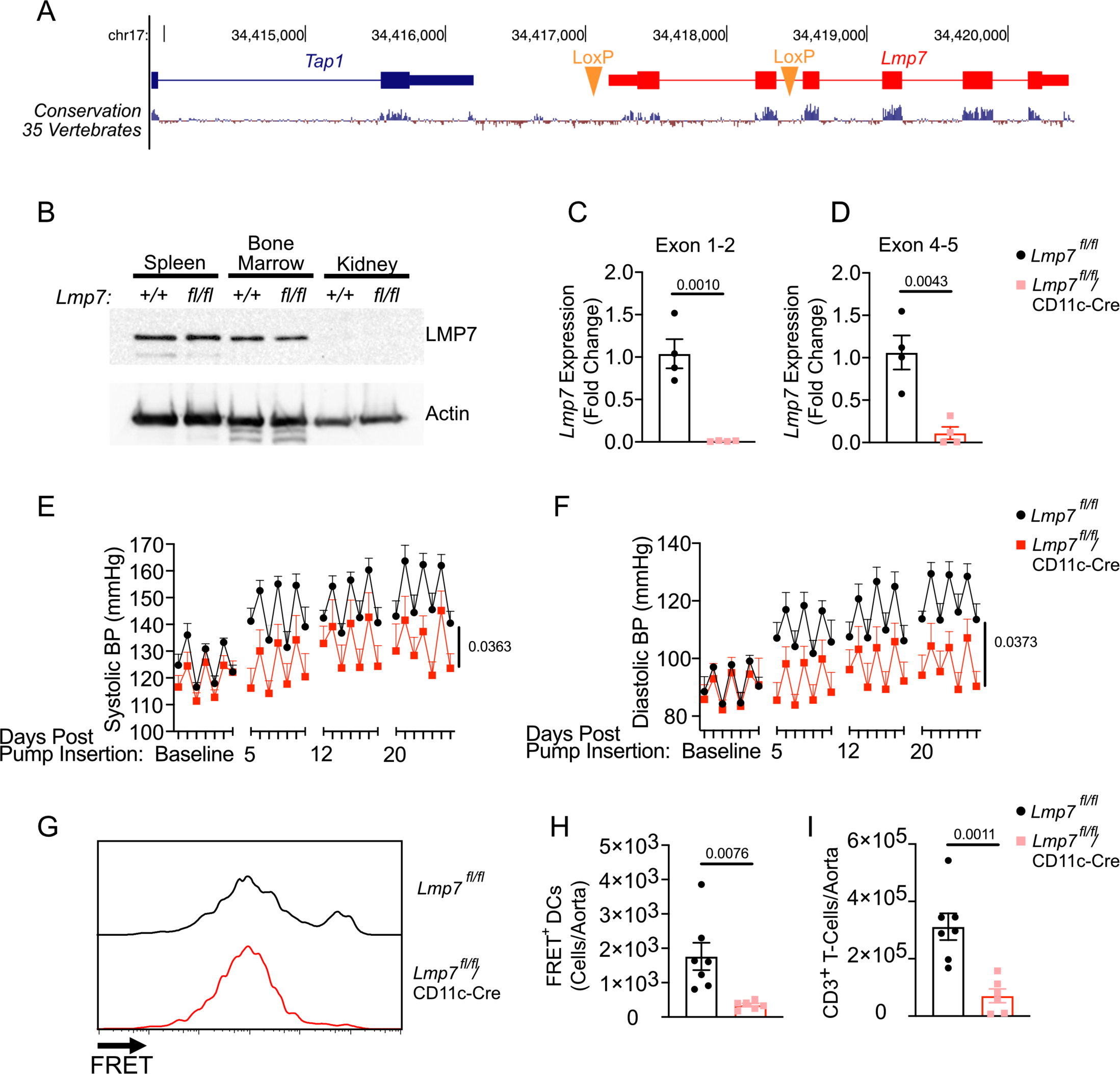
DC-specific *Lmp7* expression contributes to hypertension. **(A)** Genomic location of LoxP sites within the murine *Lmp7* locus **(B)** Immunoblot LMP7 in wild-type and *Lpm7^fl/fl^* spleen, bone marrow, and kidney. Real-time RT-PCR analysis of *Lmp7* mRNA in sorted DCs from *Lmp7^fl/fl^* and *Lmp7^fl/fl^/CD11c-Cre* using probes that detect the boundaries of **(C)** exon 1-2 and **(D)** exon 4-5 Data were analyzed by Student’s T-test or Mann-Whitney U (*n* = 4). Radiotelemetry was used to measure **(E)** systolic and **(F)** diastolic blood pressure. Blood pressure was analyzed using mixed-effects analysis (*n* = 5-6). **(G)** Representative histogram of isoLG-adduct MHC-I FRET in DCs from angiotensin II treated *Lmp7^fl/fl^* and *Lmp7^fl/fl^/CD11c-Cre* mice. **(H)** Quantitation of FRET^+^ DCs harvested from aorta. (I) Quantitation of CD3^+^ T cells harvested from aorta. Data were analyzed by two-tailed Student’s T-test or Mann-Whitney U Test (*n* = 6-7).

### ROS Production and Hypertension Increase EC-specific isoLG-adduct MHC-I Interaction and CD8^+^ T Cell Proliferation

We exposed mouse aortic endothelial cells (MAECs) to glucose oxidase (GO), an enzyme that catalyzes the formation of hydrogen peroxide from glucose in the culture medium. This was chosen to induce oxidative stress in ECs as glucose oxidase produces a more prolonged exposure to a lower concentration of ROS. Exposure of MAECs to GO resulted in an increase in ICAM-1 and I-Ab expression (Supplementary Figure 3A, 3B). Moreover, GO induced the formation of intracellular isoLG-adducts and led to augmented isoLG-adduct-H-2D^b^ FRET (Supplementary Figure 3C, 3D). ECs isolated from the aorta of Ang II-treated mice demonstrated an increase in isoLG-adduct-H-2D^b^ FRET compared to sham, confirming the presentation of isoLG adducts in the context of MHC-I in hypertensive EC’s *in vivo* (Supplementary Figure 3E, 3F). Finally, CD8^+^ T cells isolated from hypertensive mice exhibited increased proliferation when co-cultured with GO treated ECs. We did not observe increased proliferation of CD8^+^ T cells isolated from sham treated mice, nor did we observe proliferation of CD8^+^ T cells isolated from hypertensive mice when exposed to control treated ECs (Supplementary Figure 3G, 3H).

### Loss of function of LMP7 in ECs Attenuates Hypertension and Inflammation

Next we examined the role of EC-specific LMP7 expression on hypertension and associated inflammation. Immunoblot confirmed that *Lmp7^fl/fl^* mice crossed with mice expressing the EC-specific *VECAD-Cre* exhibited loss of LMP7 expression in isolated aortic ECs (Figure 6A). Importantly, *Lmp7^fl/fl^/VECAD-Cre* mice did not exhibit a reduction in LMP7 expression in non-target organs including spleen, kidney, and heart as measured by immunoblot (Figure 6B). *Lmp7^fl/fl^/VE-Cadherin-Cre* mice exhibit attenuated hypertension following Ang II infusion when compared to *Lmp7^fl/f^* controls as measured by radiotelemetry (Figure 6C; Supplementary Figure 5A, 5B). Moreover, aortic ECs harvested from AngII-infused *Lmp7^fl/fl^/VECAD-Cre* mice exhibited a reduction in isoLG-adduct-H-2D^b^ FRET compared to AngII-infused *Lmp7^fl/f^* controls (Figure 6D, 6E). Finally, aortic T-cell and DC infiltration were reduced in *Lmp7^fl/fl^/VECAD-Cre* mice compared to *Lmp7^fl/f^* controls following AngII infusion (Figure 6F-I).

**Figure 6:**
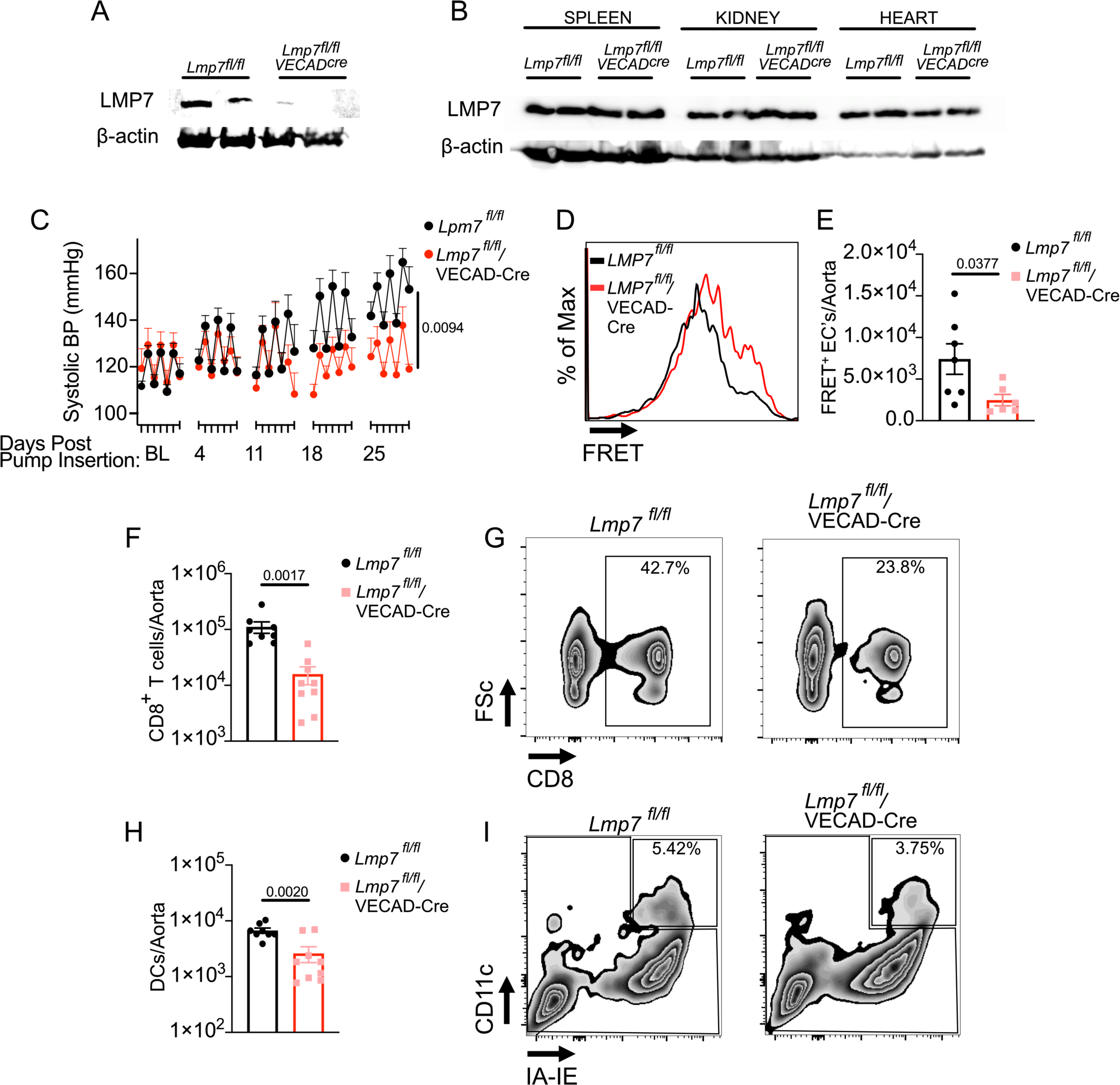
EC-specific LMP7 expression contributes to hypertension and aortic inflammation. **(A)** Immunoblot for LMP7 and β-Actin on protein isolated from ECs sorted from the aorta of *LMP7^fl/fl^* and *LMP7^fl/fl^/VECAD-Cre* mice. **(B)** Immunoblot for LMP7 and β-Actin on protein isolated from spleen, kidney, and heart of *LMP7^fl/fl^* and *LMP7^fl/fl^/VECAD-Cre* mice. Each lane represents an individual mouse. **(C)** Radiotelemetry tracings of systolic blood pressure from *LMP7^fl/fl^* and *LMP7^fl/fl^/VECAD-Cre* mice trated with Ang II. Data was analyzed by 2-way ANOVA (*n* = 5) **(D)** Quantitation and **(E)** representative histogram of FRET^+^ ECs from aorta of *LMP7^fl/fl^* and *LMP7^fl/fl^/VECAD-Cre* mice treated with Ang II. Data were analyzed by two-tailed student’s T test (*n* = 6-7). **(F)** Quantitation and **(G)** representative flow plots of infiltrating CD8^+^ T cells from aorta of *LMP7^fl/fl^* and *LMP7^fl/fl^/VECAD-Cre* mice trated with Ang II. **(H)** Quantitation and **(I)** representative flow plots of infiltrating DCs from aorta of *LMP7^fl/fl^* and *LMP7^fl/fl^/VECAD-Cre* mice treated with Ang II. Data was analyzed by two-tailed Student’s T test (*n* = 7-9).

### Scavenging of isoLGs attenuates inflammatory gene expression and STING phosphorylation in ECs

To evaluate the effects of isoLG scavengers on EC gene expression, we performed bulk RNA sequencing on GO-treated MAECs compared to MAECs co-treated with GO and the isoLG scavenger ethyl-2-hydroxybenzyloamine (Et2HOBA) as previously described.^23, 24^ Gene ontology analysis revealed downregulation of gene sets involved in the response to interferon-alpha and interferon-beta in addition to a reduction in genes involved in antigen processing and presentation (Figure 7A). Immunoblot confirmed an increase in LMP7 and H2-D^b^ expression following GO treatment. Et2HOBA co-treatment resulted in a reduction in expression of LMP7 and H2-D^b^ (Figure 7B). These findings are consistent with the activation of intracellular innate immune receptor. Importantly, previous studies have shown that proteins adducted with gamma-ketoaldehydes similar to isoLGs exhibit structural similarity to double-stranded DNA (dsDNA).^25, 26^ We therefore adducted bovine serum albumin (BSA) with increasing concentrations of isoLG and observed increased intensity of staining with the dsDNA dye Sybr Green. Adduction with 1.0 mM isoLG resulted in protein crosslinking evidenced by reduced protein mobility during electrophoresis (Figure 7C). Given the importance of the cGAS-STING pathway in the detection of intracellular dsDNA and the activation of the type-I interferon response, we examined STING phosphorylation at the activating residue serine 365._27_ GO treatment of ECs results in augmented phosphorylation of serine 365 on STING. Scavenging of isoLG attenuated STING phosphorylation (Figure 7D).

**Figure 7:**
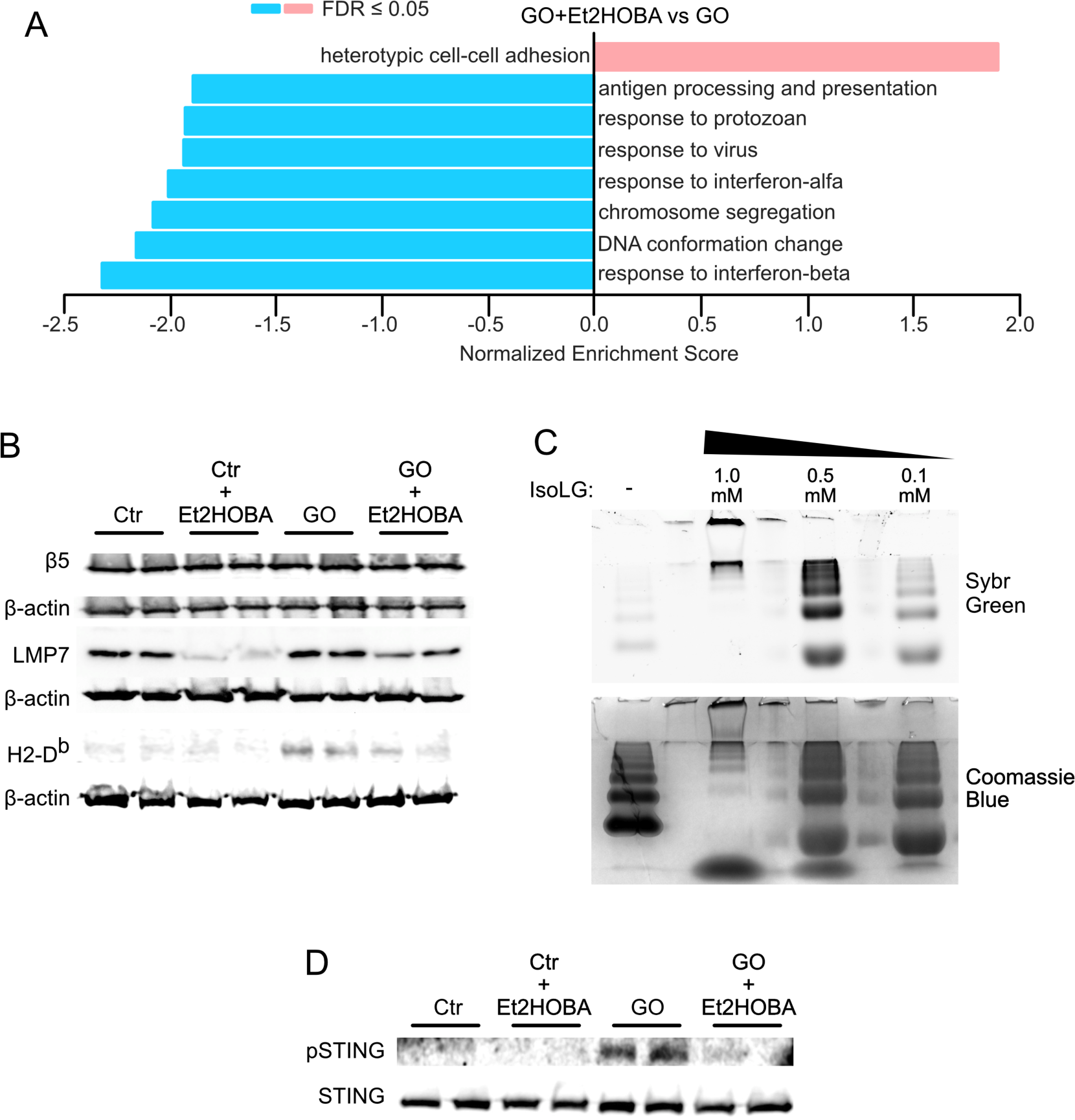
IsoLGs regulate inflammatory gene expression in ECs and contribute to STING activation. **(A)** Gene set enrichment analysis performed on sequenced RNA harvested from MAECs treated with GO or GO+Et2HOBA. **(B)** Immunoblot analysis for β5, LMP7, and H2-D^b^ with β-actin loading controls from MAECs treated with control, control+Et2HOBA, GO, or GO+Et2HOBA. **(C)** Sybr green and Coomassie blue staining of isoLG adducted BSA at increasing concentrations **(D)** Immunoblot for phospho-STING^ser365^ (pSTING) and total STING in MAECS treated with control, control+Et2HOBA, GO, or GO+Et2HOBA.

## DISCUSSION

Hypertension is the most common modifiable risk factor for the development of cardiovascular disease. It remains the primary cause of approximately 50% of strokes and ischemic myocardial events worldwide.^28^ Despite an abundance of anti-hypertensive medications, many hypertensive subjects are resistant to current therapy, and have increased risk of end-organ damage even if blood pressure is controlled. Moreover, the mechanisms underlying its development remain poorly understood. Immune activation and inflammation are clear drivers of the establishment and maintenance of hypertension. Prior work has strongly suggested that post-translational modification by isoLGs causes them to be antigenic, however how these are processed and presented to T cells has remained undefined. In the current study we show that isoLG adducts are presented by MHC-I on dendritic cells and endothelial cells by an immunoproteasome dependent mechanism. This process is required for immune activation, vascular infiltration, and blood pressure elevation in the Ang II murine model of hypertension.

In this study, we employed FRET to detect an interaction of isoLG-adducts with MHC-I. FRET occurs when fluorophores are within 10-100 Å and might have been observed if MHC-I per se were isoLG adducted. This however is unlikely as we have previously shown that MHC-I is not isoLG adducted in experimental hypertension using immunoblotting.^6^ Our finding that isoLG-adduct-H-2D^b^ FRET is virtually eliminated by proteasomal blockade further supports this conclusion. Proteasome function is critical for the processing and loading of peptides into MHC-I. Specifically, the absence of surface isoLG and isoLG-adduct-H-2D^b^ FRET with proteasome inhibition suggest a critical role of proteasome function in the establishment of this interaction.

Our studies show that isoLG-adduct-H-2D^b^ FRET is almost exclusively observed on cells expressing the surface marker CD11c in hypertensive mice. Given that CD11c is a marker of professional antigen presenting cells, it is likely that, *in vivo*, these cells are capable of isoLG adduct presentation by MHC-I and therefore play a critical role in the activation of CD8^+^ cytotoxic T cells. We have previously shown that CD11c^+^ cells produce large amounts of ROS in the setting of hypertension, and thus are likely to form isoLG adducts intracellularly.

Given the importance of the proteasome in antigen processing and presentation in immune cells, it is notable that proteasome function is augmented in DCs from hypertensive animals. Immunoproteasome function can be induced by IFNγ, and prior studies from our group and others have supported a role of this cytokine in hypertension. For example, Sun et al showed that the mineralocorticoid receptor plays a critical role in modulating IFNγ transcription in T cells, and T cell specific deletion of this receptor and infusion of an anti-IFNγ antibody eliminated the hypertensive response to Ang II infusion.^29^ We have shown that renal transporter activation in hypertension is blunted in IFNγ-deficient mice.^30^ Similarly, we have shown that IFNγ-deficient mice are protected against hypertension and renal inflation of T cells in response to repeated hypertensive stimuli.^31^ Thus, our finding of increased expression of the immunoproteasome subunit LMP7 and increased proteasome activity in hypertension might be related to the proinflammatory milieu caused by cytokines like IFNγ.^30^ These findings are consistent with the known functions of the immunoproteasome. Specifically, the immunoproteasome exhibits augmented efficiency and activity which provides a robust capacity for antigen presentation by professional antigen presenting cells during times of systemic inflammation. BTZ is a non-selective inhibitor of both constitutive and immunoproteasome function. Consistent with previous studies showing a protective effect of BTZ on aortic remodeling and cardiac hypertrophy in response to Ang II, BTZ treatment attenuated the development of hypertension in mice, however, these studies do not clearly define the relative contributions of constitutive proteasome versus immunoproteasome functions.

Previous studies have revealed that isoLGs disrupt the function of the traditional proteasome with greatest inhibition of the chymotrypsin subunit.^16^ Additionally, our findings in B8 cells support these conclusions revealing that, in the absence of immunoproteasome expression, MHC-I-isoLG-adduct interaction is low and unchanged in the presence of tBHP. Conversely, in the presence of immunoproteasome subunits, the isoLG/MHC-I interaction is robustly increased and is augmented by exposure to tBHP. This increase in interaction further supports an important role of the immunoproteasome in processing isoLG adducts for presentation in MHC-I.

We found that mice lacking all subunits of the immunoproteasome exhibit attenuated hypertension and a reduction in renal inflammation. Moreover, we found a reduction in isoLG-adduct-H-2D^b^ FRET in DCs associated with the aorta. Specific inhibition of the chymotrypsin immunoproteasome subunit LMP7 with PR-957 disrupts isoLG-adduct-H-2D^b^ FRET in aortic DCs and reduces T cell infiltration into the aorta. PR-957 treatment phenocopies the attenuation of blood pressure observed in mice co-treated with Ang II and BTZ and in TKO mice treated with Ang II. Conditional deletion of *LMP7* in DCs and ECs results in attenuation of hypertension and aortic T-cell infiltration in response to Ang II. This is associated with a reduction in IsoLG-adduct-H-2D^b^ FRET. This suggests that the effects observed with BTZ and in the TKO animals is primarily driven by its effects on LMP7.

Following activation, it has been established that CD8^+^ T cells migrate into peripheral tissues including the vascular wall in hypertension.^2^ It has been suggested that homing of T cells is guided by display of similar antigens on MHC-I by the endothelium. In the current study we show that, similar to DCs, ECs also present isoLG-adducts within the context of MHC-I by an LMP7-dependent mechanism. Display of these adducts in MHC-I results in T-cell proliferation, migration, and activation. ROS production within ECs is a hallmark of hypertension. ECs express several Nox subunits of the NADPH oxidase, capable of producing ROS in response to numerous stimuli including mechanical forces and inflammatory cytokines.^32, 33^ Moreover, nitric oxide synthase uncoupling in ECs may result from numerous stimuli including ROS produced by Nox2, leading to production of superoxide instead of nitric oxide (NO).^34^ These ROS may contribute to the generation of intracellular isoLG-adducts in ECs leading to isoLG-adduct display in MHC-I. These studies thus defined a previously unknown mechanism by which T cell homing to target organs occur.

The regulation of LMP7 expression by isoLGs is intriguing. This is accompanied by a strong reduction in gene programs involved in response to interferons in ECs treated with an isoLG scavenger. Previous work has revealed a critical role of type I and type II interferons in the regulation of LMP7.^35, 36^ Taken together, these findings support a role of isoLGs as direct activators of interferon response genes. Previous studies have revealed that proteins adducted with 4-oxo-2-nonenol (ONE), a γ-ketoaledhyde similar in structure to isoLGs, bind with DNA intercalating dye in a dose-dependent manner and cross-react with anti-DNA antibodies.^26^ We found that BSA adducted with isoLG interacts with a dsDNA-specific dye suggesting a structural similarity of isoLG adducts with dsDNA. The isoLG-dependence of STING phosphorylation in ECs suggests that the DNA mimicry of isoLG adducts leads to activation of cGAS. STING is a crucial part of the cyclic GMP-AMP synthase (cGAS) pathway, which recognizes viral dsDNA in the cytosol.^27^ Taken together with the finding that the immunoproteasome is more efficient at isoLG processing than the constitutive proteasome, these data suggest a novel mechanism whereby isoLG adduct accumulation activates the expression of immunoproteasome subunits via the cGAS-STING pathway leading to the enhanced clearance of isoLG adducts and presentation within MHC-I. Future studies will focus on the role of isoLG adducts as activators of dsDNA-responsive innate immune receptors in ECs and DCs.

This study further confirms an important role of isoLG-adducted antigens as drivers of T cell proliferation, migration, and activation in hypertension. This discovery is remarkable and therapeutically important given the implied potential for immunotherapies upon discovery of unique antigens. This study also confirms that MHC-I molecules are direct mediators of hypertension.

Numerous GWAS analyses have been performed but have not identified unique HLA haplotypes associated with hypertension, however, in depth haplotyping of MHC-I alleles has not been performed in a hypertensive population. It is likely that unique MHC-I alleles predispose individuals to the development of hypertension and that this predisposition may be directly related to the capacity of each allele to display isoLG adducted antigens. These will be the topics of future studies.

## STAR*Methods

## RESOURCE AVAILABILITY

### Lead Contact

Further information and requests for resources and reagents should be directed to and will be fulfilled by the lead contact David Patrick.

### Materials availability

Upon specific reasonable request and execution of a material transfer agreement (MTA) from Vanderbilt University Medical Center to the lead contact, reagents will be made available.

### Data and code availability

RNA sequencing data have been deposited at GEO and are publicly available as of the date of publication. Accession numbers are listed in the key resource table. Original western blot images are publicly available as of the date of this publication. All data reported in this paper will be shared by the lead contact upon request.

## EXPERIMENTAL MODEL AND STUDY PARTICIPANT DETAILS

### Mice

*C57BL/6J* mice were obtained from The Jackson Laboratory. Immunoproteasome triple knockout (TKO) mice were obtained from Regeneron Therapeutics. *Lmp7^fl/fl^* were generated on the *C57BL/6* background by Gene Edit Biolab (Atlanta, GA, USA). *CD11c-Cre* and *VE-Cadherin-Cre* (*VECAD-Cre*) mice were obtained from Jackson Laboratory. Immunoproteasome triple knockout mice (TKO) were obtained from Regeneron Therapeutics. Stock numbers for commercially available strains are listed in the key resources table. All animals studied were male which limits the generalizability of the study. All animal procedures were approved by Vanderbilt University Medical Center’s Institutional Animal Care and Use Committee, and the mice were housed and cared for in accordance with the Guide for the Care and Use of Laboratory Animals, US Department of Health and Human Services.

### Cell lines

B8 and B8.27M.2 cells were a gift from Marcus Groettrup and have been published previously. ^21^ Cells were maintained in IMDM medium (IMDM supplemented with 10% FBS, 2 mM L-Glutamine, 50 µM beta-mercaptoethanol. Cells were kept in a 5% carbon dioxide (CO_2_) incubator at 37°C. C57BL/6 mouse primary aortic endothelial cells (MAECs) were obtained from Cell Biologics (C57-6052) and were incubated in Complete Endothelial Cell Medium (Cell Biologics, M1168) in a 5% carbon dioxide (CO2) incubator at 37°C.

### Human Samples

Subjects with hypertension and controls were recruited by advertisement to the Vanderbilt Clinical Trials Center as part of the Immune Mechanisms of Essential and Lupus-Related Hypertension study. These subjects were 31-56 years old. Exclusion criteria included confirmed or suspected renal, renovascular, or endocrine causes of secondary hypertension, concomitant diabetes, concomitant illness requiring corticosteroids or immunosuppressants, recent (within 3-months) vaccination against any infectious agent, active ongoing malignancy, severe psychiatric disorders, or HIV/AIDS. All subjects gave written informed consent. Patient demographics including age, race, sex, and blood pressure are presented in Supplemental Table 1. The institutional review board of Vanderbilt University Medical Center approved the human study (IRB# 150544).

## METHOD DETAILS

### BTZ and PR-957 Treatment of Mice

Animals were treated with Ang II and blood pressure was measured by radiotelemetry as previously described.^6^ BTZ (LC Labs) was dissolved in saline and injected at 0.5 mg/kg by an intraperitoneal route three times per week. PR-957 was dissolved in 10% betadex sulfobutyl ether (sigma) and 10 mM sodium citrate (pH 3.5) and administered intraperitoneally to mice at 6 mg/kg three times per week. Blood pressure was monitored noninvasively using tail cuff and invasively using radiotelemetry as previously described.^6, 37^

### Flow cytometry

Tissue homogenates from mice were filtered through a 40 μm filter. Single cell suspensions were stained for flow cytometry and analyzed in separate panels using the antibodies and fluorophores shown in Supplemental Table 1. A known quantity of calibration (counting) beads was added to each sample before analysis. Samples were run on a BD FACSCanto II system or a Cytek Aurora system and analyzed using FloJo software. Gates were set using fluorescence minus one controls. FRET was performed on the BD FACSCanto II using PE and AF-647 as a donor acceptor pair. FRET was performed on the Cytek Aurora using BV605 and APC/Cy7 as a donor acceptor pair. Staining for isoLG adducts was performed using the single chain antibody D11, which recognized isoLG lysines independent of adjacent amino acids. The D11 antibody was labeled with a fluorochrome using the APEX Alexa Fluor 488 or APEX Alexa Fluor 647 Antibody Labeling Kit (ThermoFisher Scientific). D11 was labeled with Biotin using a biotinylation kit (Abcam). Additionally, activation of the immune response was observed in endothelial cells staining for ICAM and MHC II (IA/IE). Vascular immune infiltration was studied through the populations of macrophages (CD64^+^, MERTK^+^), dendritic cells (CD11c^+^, IA/IE^+^), B cells (CD19^+^), cytotoxic T cells (CD3^+^, CD8^+^) and CD4^+^ T cells (CD3^+^, CD4^+^).

### TBHP and glucose oxidase treatment of cells

All tBHP treatments were performed at a final concentration of 1 mM tBHP for 30-minutes. CD11c^+^ cells were sorted from spleens of Sham and Ang II treated *C57Bl/6* mice using a commercially available kit following the manufacturer’s instructions (Miltenyi). Cells were pre-treated with 10 μM MG-132 (Cell Signaling) or 10 nM or 100 nM BTZ (Cell Signaling) for 1 hour and then co-treated with 100 μM tBHP for 30-minutes. Medium was then changed to remove tBHP and cells were treated for an additional 24 hours with aforementioned proteasome inhibitor. Proteasome activity assays were performed using the ProteasomeGlo assay following the manufacturer’s instructions (BioRad). To model endothelial cell ROS production, we incubated mouse aortic endothelial cells (MAECs, Cell biologics) with glucose and glucose oxidase (GO, 5 mU/mL) for 2 hours and then changed medium following this stimulus. In additional experiments, groups treated with the specific scavenger for IsoLG adducts ethyl-2-hydroxybenzylamine (Et2HOBA, 0.2 mM) were included. The cells were collected for testing 24h after addition of the original stimulus.

### IsoLG Adduction, electrophoresis, and staining

50 μg of bovine serum albumin (BSA) was dissolved in phosphate buffered saline (PBS) at a concentration of 1 mg/mL. Un-reacted isoLG was synthesized as previously described.^38^ BSA was adducted at 0.1, 0.5, and 1 mM final concentration of isoLG at 4°C overnight. 6x non-denaturing loading dye (0.03% bromophenol blue, 60% glycerol, 1x TBE) was added to each sample to a final concentration of 1x. Samples were separated using non-denaturing polyacrylamide gel electrophoresis. Gels were then incubated with Bio-Safe Coomassie Stain following the manufacturer’s instructions (BioRad) or in 20 mL TBE + 5 uL of SYBR Green DNA Gel Stain (ThermoFisher Scientific). Gels were imaged on the BioRad Gel Doc imaging system.

### Human monocyte isolation

Peripheral blood mononuclear cells were prepared using anti-CD14 magnetic microbeads and sorted (Miltenyi) as previously described. ^6, 39^

### Immunoblot

For immunoblot, cells were lysed with RIPA buffer and 50 μg of protein was separated by SDS-PAGE. Gels were transferred to a nitrocellulose membrane and incubated with 1:1000 of the anti-human-LMP7 antibody or anti-mouse/human/rat LMP7 antibody, phospho-STING^ser365^ (S365, 1:1000), STING (1:1000), MHC-I H2 Db/D1 (1:200). The blot was stripped with Restore Western Blot Stripping Buffer (ThermoFisher Scientific) and reprobed with anti-β-actin-Peroxidase antibody clone AC-15 (Sigma) at a 1:20,000 dilution or anti-GAPDH from Abcam at a 1:5000 dilution. Imaging was performed with autoradiography film or a BioRad VersaDock. Western blots were quantitated with ImageJ (NIH).

### T cell proliferation studies

To determine if endothelial ROS production affects T cell proliferation, we exposed MAECs to hydrogen peroxide generated by glucose and GO for 2 h. T-cells were harvested by magnetic sorting from the bone marrow mice treated for two weeks with sham or 490 ng/kg/min Ang II as previously described.^6^ After an additional 24 hours, medium was changed to remove GO and 0.5 x 10^6^ T-cells from HLA-matched *C57Bl/6* mice were added to the culture dish. Prior to incubation, cells were loaded with the cell proliferation marker CFSE. T-cells and MEACs were co-cultured for 96 hours. Proliferation was determined using flow cytometry.

### Bulk RNA sequencing

We incubated MAECs with glucose oxidase and collected the RNA 24h after the stimulus using the RNeasy Mini Kit from QIAGEN. Four samples per experimental group (control, glucose oxidase, and glucose oxidase + Et2HOBA) were pooled and next generation sequencing was performed. Data was analyzed and the most affected biological processes were identified through Web Gestalt using geneontology as the functional database for Gene Set Enrichment Analysis (GSEA).^40^ Sequencing was performed at Paired-End 150 bp on the Illumina NovaSeq 6000 targeting an average of 50M reads per sample. An average of 50M reads per sample was targeted. The data was aligned to mm10 genome by STAR softwatre, and then the reads were extracted by FeatureCount software. Differential detection was performed byDESeq2 software.

## QUANTIFICATION AND STATISTICAL ANALYSIS

All data are expressed as mean ± SEM. Comparisons made between 2 variables were performed using Student’s t tests or Mann-Whitney U test depending on normality of distribution. Normality of distribution of data was confirmed using the D’Agostino-Pearson normality test. Comparisons among more than 2-variables were performed with 1-way ANOVA with Tukey’s post-hoc test. To compare differences in blood pressure and proteasome activity, 2-way ANOVA followed by Šídák’s post-hoc test was used. Mixed effects analysis was used to compare blood pressure measurements for which there is missing data. Clinical parameters were compared with 2-way ANOVA followed by Bonferroni post-hoc test. Differences were considered significant at p < 0.05. Data were analyzed using GraphPad Prism v9 (GraphPad Software) or software indicated in relevant methods section.

## Supporting information

Key Resources Table

## ACKNOWLEDGEMENTS

This work was supported by National Institute of Health grants 5R35HL140016-02, 5P01HL129941-03, 1R01HL134895-02. Additional funding support was provided by the Veterans Affairs Biomedical Laboratory Research Career Development Award IK2BX005376. Dr. de la Visitación was supported by an American Heart Association post-doctoral fellowship (915301). Dr. van Beusecum was supported by Veterans Affairs Biomedical Laboratory Research Career Development Award 1IK2BX005605-01. We would also like to acknowledge Marcus Groettrup, University of Konstanz, for the generous gift of the B8 and B8.27M.2 cells.

## AUTHOR CONTRIBUTIONS

NV and DMP designed the study. NV, WC, JK, JPVB, EMH, MS, MA, and DMP performed experiments and acquired the data. NV, DMP, DGH, JPVB, and JK analyzed the data. SZ assisted with RNA sequencing analysis. VA synthesized and prepared the Et2HOBA. NV and DMP wrote the manuscript. All authors reviewed and revised the manuscript. DMP supervised the study. *

## REFERENCES

1. Sesso HD, Buring JE, Rifai N, Blake GJ, Gaziano JM and Ridker PM. C-reactive protein and the risk of developing hypertension. JAMA. 2003;290:2945–51.

2. Itani HA, McMaster WG, Jr., Saleh MA, Nazarewicz RR, Mikolajczyk TP, Kaszuba AM, Konior A, Prejbisz A, Januszewicz A, Norlander AE, Chen W, Bonami RH, Marshall AF, Poffenberger G, Weyand CM, Madhur MS, Moore DJ, Harrison DG and Guzik TJ. Activation of Human T Cells in Hypertension: Studies of Humanized Mice and Hypertensive Humans. Hypertension. 2016;68:123–32.

3. Mikolajczyk TP, Nosalski R, Szczepaniak P, Budzyn K, Osmenda G, Skiba D, Sagan A, Wu J, Vinh A, Marvar PJ, Guzik B, Podolec J, Drummond G, Lob HE, Harrison DG and Guzik TJ. Role of chemokine RANTES in the regulation of perivascular inflammation, T-cell accumulation, and vascular dysfunction in hypertension. FASEB J. 2016;30:1987–99.

4. Youn JC, Yu HT, Lim BJ, Koh MJ, Lee J, Chang DY, Choi YS, Lee SH, Kang SM, Jang Y, Yoo OJ, Shin EC and Park S. Immunosenescent CD8+ T cells and C-X-C chemokine receptor type 3 chemokines are increased in human hypertension. Hypertension. 2013;62:126–33.

5. Liu Y, Rafferty TM, Rhee SW, Webber JS, Song L, Ko B, Hoover RS, He B and Mu S. CD8(+) T cells stimulate Na-Cl co-transporter NCC in distal convoluted tubules leading to salt-sensitive hypertension. Nat Commun. 2017;8:14037.

6. Kirabo A, Fontana V, de Faria AP, Loperena R, Galindo CL, Wu J, Bikineyeva AT, Dikalov S, Xiao L, Chen W, Saleh MA, Trott DW, Itani HA, Vinh A, Amarnath V, Amarnath K, Guzik TJ, Bernstein KE, Shen XZ, Shyr Y, Chen SC, Mernaugh RL, Laffer CL, Elijovich F, Davies SS, Moreno H, Madhur MS, Roberts J and Harrison DG. DC isoketal-modified proteins activate T cells and promote hypertension. J Clin Invest. 2014;124:4642–56.

7. Davies SS, May-Zhang LS, Boutaud O, Amarnath V, Kirabo A and Harrison DG. Isolevuglandins as mediators of disease and the development of dicarbonyl scavengers as pharmaceutical interventions. Pharmacol Ther. 2020;205:107418.

8. Lozanoska-Ochser B and Peakman M. Level of major histocompatibility complex class I expression on endothelium in non-obese diabetic mice influences CD8 T cell adhesion and migration. Clin Exp Immunol. 2009;157:119–27.

9. Pober JS, Merola J, Liu R and Manes TD. Antigen Presentation by Vascular Cells. Front Immunol. 2017;8:1907.

10. Pober JS. Immunobiology of human vascular endothelium. Immunol Res. 1999;19:225–32.

11. Rousseau A and Bertolotti A. Regulation of proteasome assembly and activity in health and disease. Nat Rev Mol Cell Biol. 2018;19:697–712.

12. Kloetzel PM. Antigen processing by the proteasome. Nat Rev Mol Cell Biol. 2001;2:179–87.

13. Dick TP, Nussbaum AK, Deeg M, Heinemeyer W, Groll M, Schirle M, Keilholz W, Stevanovic S, Wolf DH, Huber R, Rammensee HG and Schild H. Contribution of proteasomal beta-subunits to the cleavage of peptide substrates analyzed with yeast mutants. J Biol Chem. 1998;273:25637–46.

14. Mishto M, Liepe J, Textoris-Taube K, Keller C, Henklein P, Weberruss M, Dahlmann B, Enenkel C, Voigt A, Kuckelkorn U, Stumpf MP and Kloetzel PM. Proteasome isoforms exhibit only quantitative differences in cleavage and epitope generation. Eur J Immunol. 2014;44:3508–21.

15. Gaczynska M, Rock KL, Spies T and Goldberg AL. Peptidase activities of proteasomes are differentially regulated by the major histocompatibility complex-encoded genes for LMP2 and LMP7. Proc Natl Acad Sci U S A. 1994;91:9213–7.

16. Davies SS, Amarnath V, Montine KS, Bernoud-Hubac N, Boutaud O, Montine TJ and Roberts LJ, 2nd. Effects of reactive gamma-ketoaldehydes formed by the isoprostane pathway (isoketals) and cyclooxygenase pathway (levuglandins) on proteasome function. FASEB J. 2002;16:715–7.

17. Li S, Wang X, Li Y, Kost CK, Jr. and Martin DS. Bortezomib, a proteasome inhibitor, attenuates angiotensin II-induced hypertension and aortic remodeling in rats. PLoS One. 2013;8:e78564.

18. Ren H, Li F, Tian C, Nie H, Wang L, Li HH and Zheng Y. Inhibition of Proteasome Activity by Low-dose Bortezomib Attenuates Angiotensin II-induced Abdominal Aortic Aneurysm in Apo E(-/-) Mice. Sci Rep. 2015;5:15730.

19. Li J, Wang S, Zhang YL, Bai J, Lin QY, Liu RS, Yu XH and Li HH. Immunoproteasome Subunit beta5i Promotes Ang II (Angiotensin II)-Induced Atrial Fibrillation by Targeting ATRAP (Ang II Type I Receptor-Associated Protein) Degradation in Mice. Hypertension. 2019;73:92–101.

20. Xie X, Bi HL, Lai S, Zhang YL, Li N, Cao HJ, Han L, Wang HX and Li HH. The immunoproteasome catalytic beta5i subunit regulates cardiac hypertrophy by targeting the autophagy protein ATG5 for degradation. Sci Adv. 2019;5:eaau0495.

21. Groettrup M, Standera S, Stohwasser R and Kloetzel PM. The subunits MECL-1 and LMP2 are mutually required for incorporation into the 20S proteasome. Proc Natl Acad Sci U S A. 1997;94:8970–5.

22. Kincaid EZ, Che JW, York I, Escobar H, Reyes-Vargas E, Delgado JC, Welsh RM, Karow ML, Murphy AJ, Valenzuela DM, Yancopoulos GD and Rock KL. Mice completely lacking immunoproteasomes show major changes in antigen presentation. Nat Immunol. 2011;13:129–35.

23. Patrick DM, de la Visitacion N, Krishnan J, Chen W, Ormseth MJ, Stein CM, Davies SS, Amarnath V, Crofford LJ, Williams JM, Zhao S, Smart CD, Dikalov S, Dikalova A, Xiao L, Van Beusecum JP, Ao M, Fogo AB, Kirabo A and Harrison DG. Isolevuglandins disrupt PU.1-mediated C1q expression and promote autoimmunity and hypertension in systemic lupus erythematosus. JCI Insight. 2022;7.

24. Krishnan J, de la Visitacion N, Hennen EM, Amarnath V, Harrison DG and Patrick DM. IsoLGs (Isolevuglandins) Drive Neutrophil Migration in Hypertension and Are Essential for the Formation of Neutrophil Extracellular Traps. Hypertension. 2022;79:1644–1655.

25. Otaki N, Chikazawa M, Nagae R, Shimozu Y, Shibata T, Ito S, Takasaki Y, Fujii J and Uchida K. Identification of a lipid peroxidation product as the source of oxidation-specific epitopes recognized by anti-DNA autoantibodies. J Biol Chem. 2010;285:33834–42.

26. Miyashita H, Chikazawa M, Otaki N, Hioki Y, Shimozu Y, Nakashima F, Shibata T, Hagihara Y, Maruyama S, Matsumi N and Uchida K. Lysine pyrrolation is a naturally-occurring covalent modification involved in the production of DNA mimic proteins. Sci Rep. 2014;4:5343.

27. Hopfner KP and Hornung V. Molecular mechanisms and cellular functions of cGAS-STING signalling. Nat Rev Mol Cell Biol. 2020;21:501–521.

28. Lawes CM, Vander Hoorn S, Rodgers A and International Society of H. Global burden of blood-pressure-related disease, 2001. Lancet. 2008;371:1513-8.

29. Sun XN, Li C, Liu Y, Du LJ, Zeng MR, Zheng XJ, Zhang WC, Liu Y, Zhu M, Kong D, Zhou L, Lu L, Shen ZX, Yi Y, Du L, Qin M, Liu X, Hua Z, Sun S, Yin H, Zhou B, Yu Y, Zhang Z and Duan SZ. T-Cell Mineralocorticoid Receptor Controls Blood Pressure by Regulating Interferon-Gamma. Circ Res. 2017;120:1584–1597.

30. Kamat NV, Thabet SR, Xiao L, Saleh MA, Kirabo A, Madhur MS, Delpire E, Harrison DG and McDonough AA. Renal transporter activation during angiotensin-II hypertension is blunted in interferon-gamma-/- and interleukin-17A-/- mice. Hypertension. 2015;65:569–76.

31. Itani HA, Xiao L, Saleh MA, Wu J, Pilkinton MA, Dale BL, Barbaro NR, Foss JD, Kirabo A, Montaniel KR, Norlander AE, Chen W, Sato R, Navar LG, Mallal SA, Madhur MS, Bernstein KE and Harrison DG. CD70 Exacerbates Blood Pressure Elevation and Renal Damage in Response to Repeated Hypertensive Stimuli. Circ Res. 2016;118:1233–43.

32. Loperena R and Harrison DG. Oxidative Stress and Hypertensive Diseases. Med Clin North Am. 2017;101:169–193.

33. Drummond GR and Sobey CG. Endothelial NADPH oxidases: which NOX to target in vascular disease? Trends Endocrinol Metab. 2014;25:452–63.

34. Kossmann S, Hu H, Steven S, Schonfelder T, Fraccarollo D, Mikhed Y, Brahler M, Knorr M, Brandt M, Karbach SH, Becker C, Oelze M, Bauersachs J, Widder J, Munzel T, Daiber A and Wenzel P. Inflammatory monocytes determine endothelial nitric-oxide synthase uncoupling and nitro-oxidative stress induced by angiotensin II. J Biol Chem. 2014;289:27540–50.

35. Yang I, Kremen TJ, Giovannone AJ, Paik E, Odesa SK, Prins RM and Liau LM. Modulation of major histocompatibility complex Class I molecules and major histocompatibility complex-bound immunogenic peptides induced by interferon-alpha and interferon-gamma treatment of human glioblastoma multiforme. J Neurosurg. 2004;100:310–9.

36. Castellano G, Cafiero C, Divella C, Sallustio F, Gigante M, Pontrelli P, De Palma G, Rossini M, Grandaliano G and Gesualdo L. Local synthesis of interferon-alpha in lupus nephritis is associated with type I interferons signature and LMP7 induction in renal tubular epithelial cells. Arthritis Res Ther. 2015;17:72.

37. Van Beusecum JP, Barbaro NR, Smart CD, Patrick DM, Loperena R, Zhao S, de la Visitacion N, Ao M, Xiao L, Shibao CA and Harrison DG. Growth Arrest Specific-6 and Axl Coordinate Inflammation and Hypertension. Circ Res. 2021;129:975–991.

38. Amarnath V, Amarnath K, Masterson T, Davies S and Roberts LJ. A Simplified Synthesis of the Diastereomers of Levuglandin E2. Synthetic Communications. 2005;35:397–408.

39. Norlander AE, Saleh MA, Pandey AK, Itani HA, Wu J, Xiao L, Kang J, Dale BL, Goleva SB, Laroumanie F, Du L, Harrison DG and Madhur MS. A salt-sensing kinase in T lymphocytes, SGK1, drives hypertension and hypertensive end-organ damage. JCI Insight. 2017;2.

40. Liao Y, Wang J, Jaehnig EJ, Shi Z and Zhang B. WebGestalt 2019: gene set analysis toolkit with revamped UIs and APIs. Nucleic Acids Res. 2019;47:W199–W205.

