## Supplemental Material for "Immunoproteasomal Processing of Isolevuglandin Adducts in Hypertension"

**Affiliations:** ^1^Division of Clinical Pharmacology, Department of Medicine, Vanderbilt University Medical Center, Nashville, Tennessee, USA. ^2^Department of Veterans Affairs, Charleston South Carolina. ^3^Department of Biomedical Engineering, Vanderbilt University.  ^4^Vanderbilt Center for Quantitative Science, Vanderbilt University Medical Center. ^5^Division of Cardiovascular Medicine, Department of Medicine, Vanderbilt University Medical Center. ^6^ Department of Veterans Affairs, Nashville, Tennessee.

Word Count: 6,277

*Author for Correspondence

David M. Patrick MD, PhD

VA Medical Center

1310 24^th^ Avenue S.

Nashville, TN 37212

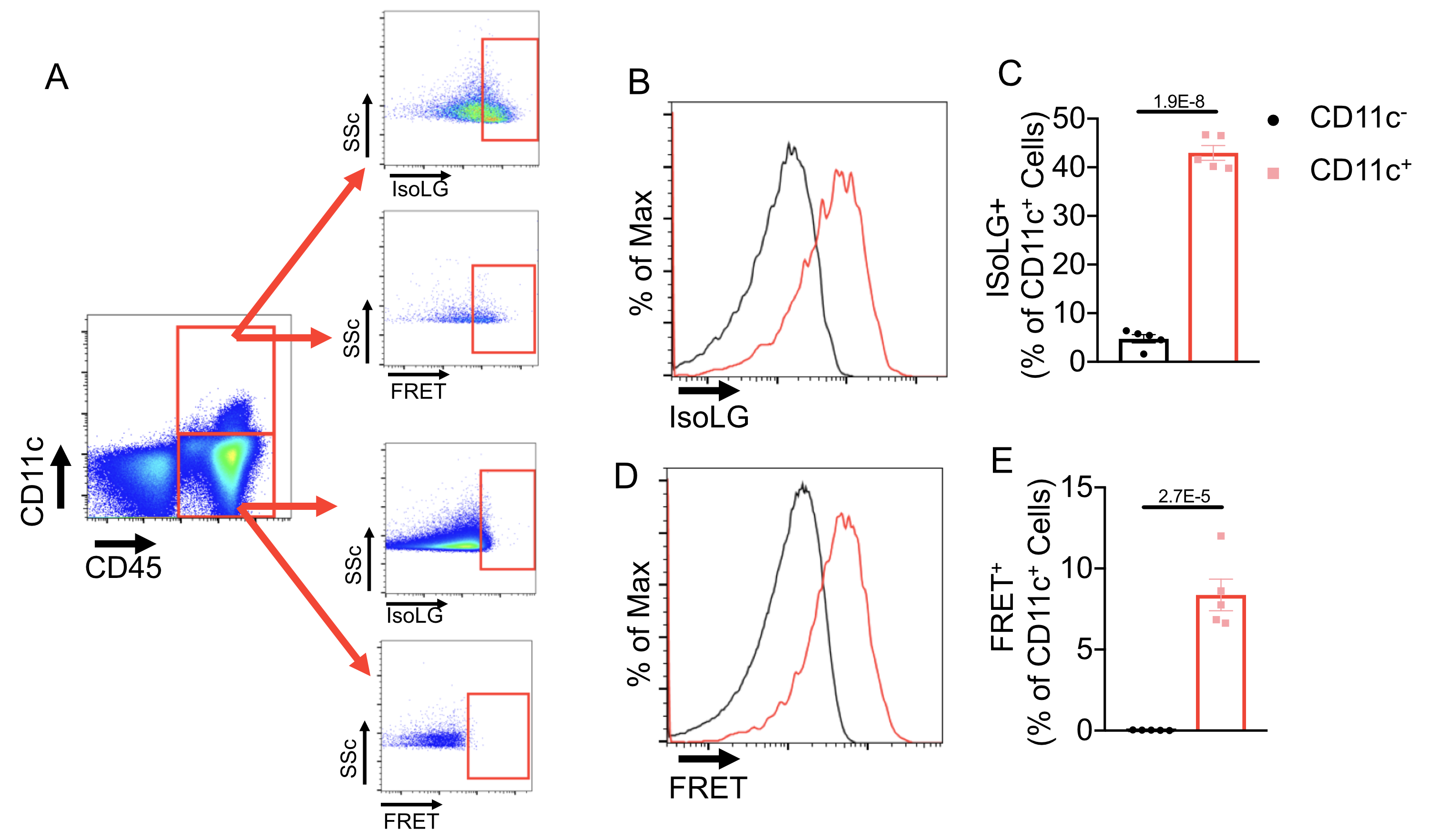

**Supplemental Figure 1. IsoLG H-2D^b^ interaction in hypertensive mice is restricted to CD11c^+^ cells. (A)** Gating strategy to determine the presence of surface isoLG and isoLG-H-2D^b^ FRET. (B) Histogram of surface isoLG staining of CD11c^-^ and CD11c^+^ cells. (C) Quantitation of isoLG^+^ CD11c^-^ and CD11c^+^ cells. (D) Histogram of FRET in CD11c^-^ and CD11c^+^ cells. (E) Quantitation of FRET in CD11c^-^ and CD11c^+^ cells. Data were analyzed by two-tailed Student’s T-test (*n* = 5).

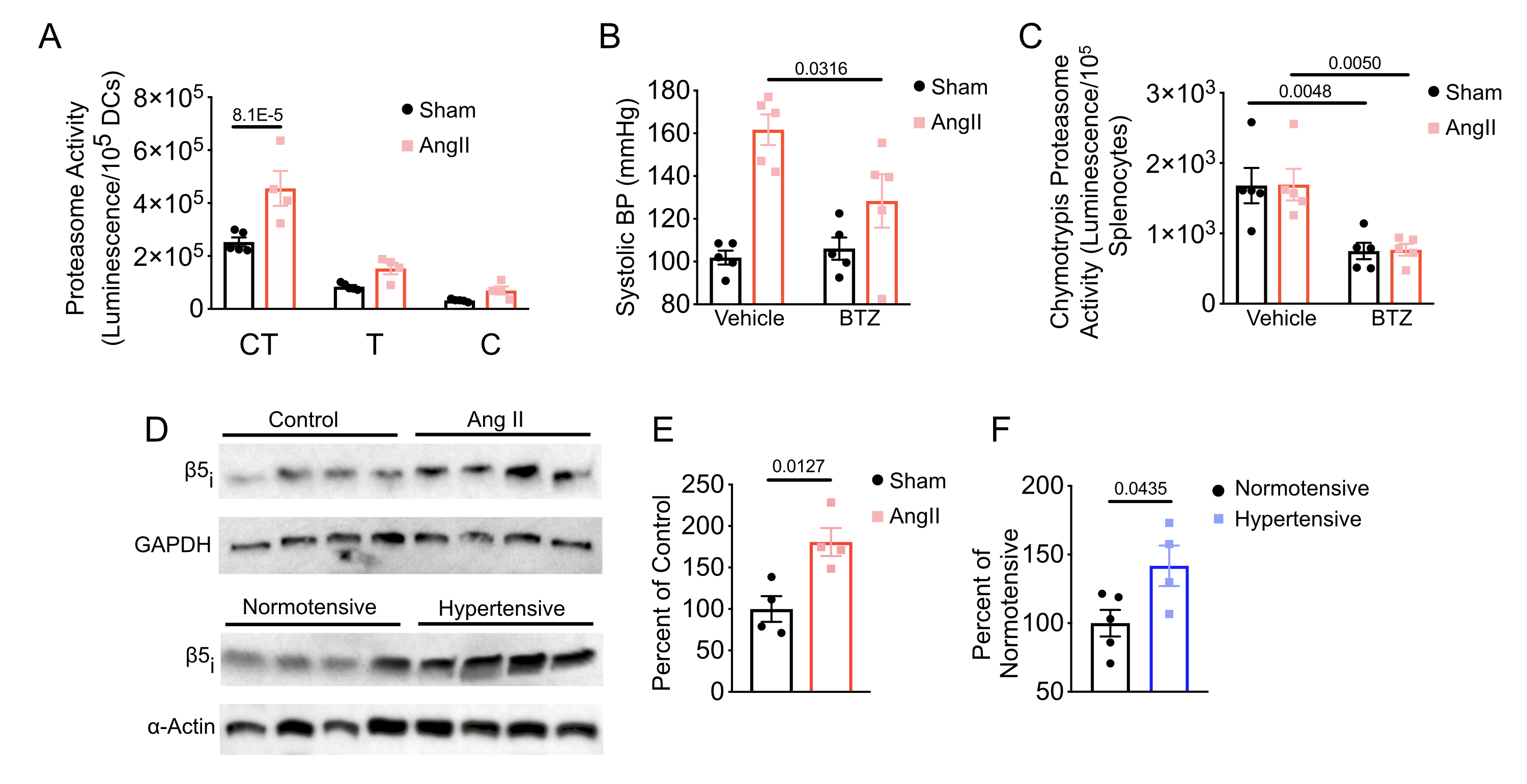

**Supplemental Figure 2. DC specific Proteasome activity is augmented in and contributes to hypertension. (A)** Chymotrypsin-like (CT), trypsin-like (T), and caspase-like (C) proteasome activity was determined in sorted CD11c^+^ splenocytes from Sham and Ang II treated mice. Data were analyzed by 2-way ANOVA with Šídák's multiple comparison test (*n* = 4-5). **(B)** Systolic blood pressure in co-treated with Sham or Ang II and Vehicle or BTZ. **(C)** Chymotrypsin proteasome activity is reduced in total splenocytes with BTZ treatment. Data were analyzed 2-way ANOVA with Šídák's multiple comparison test (*n* = 5). **(D)** Immunoproteasome chymotrypsin subunit β5_i_ expression is increased in DCs from Ang II treated mice and CD14^+^ monocytes from hypertensive patients. **(E)** Quantitation of protein levels of β5_i_ expression from DCs harvested from sham or Ang II treated mice. **(F)** Quantitation of protein levels of β5_i_ expression from CD14^+^ monocytes harvested from normotensive or hypertensive subjects. Data were analyzed by two-tailed Student’s T-test (*n* = 4-5).

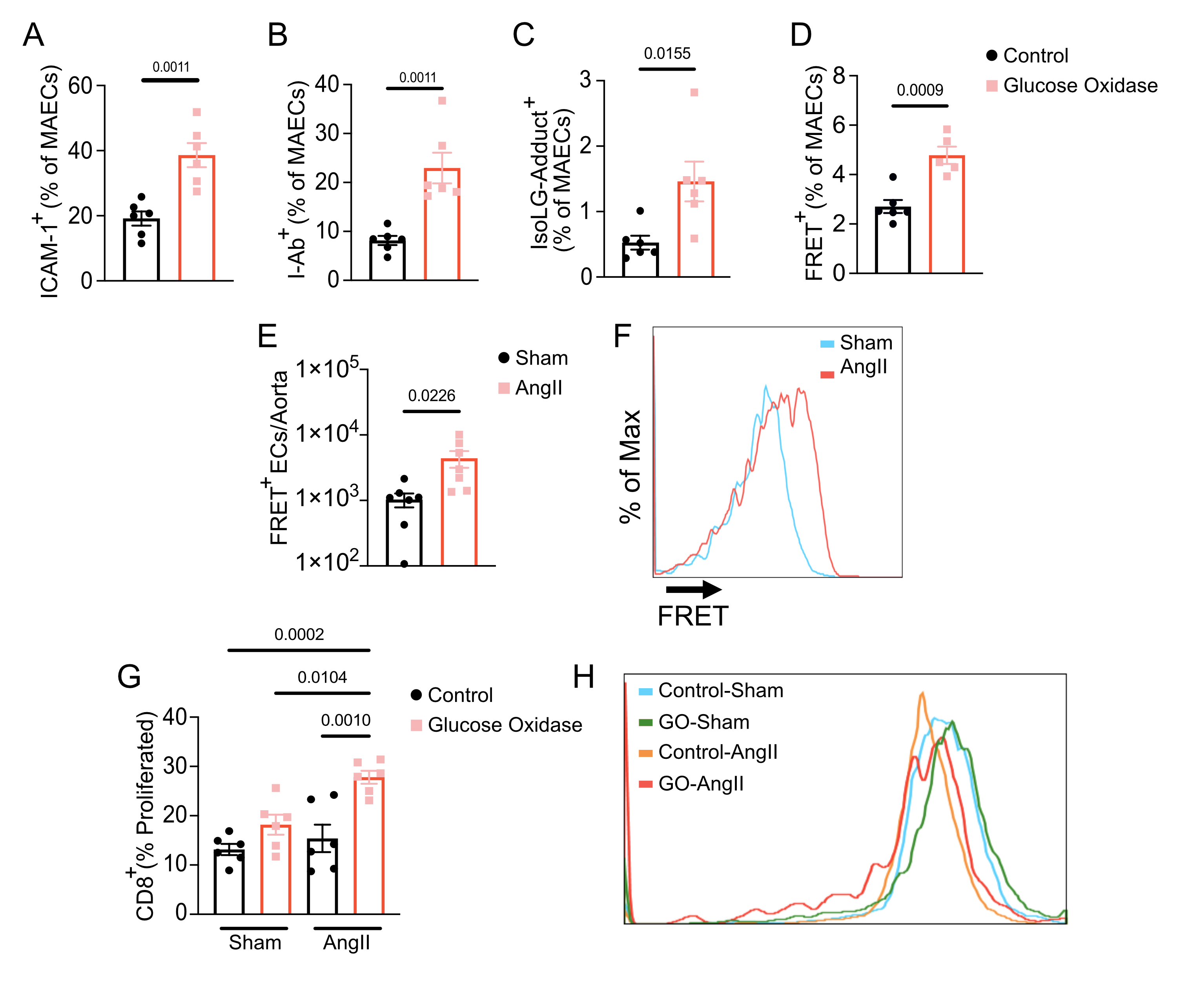

**Supplemental Figure 3. ROS Increases IsoLG-Adduct-H-2D^b^ interaction in ECs and drives T cell proliferation.** Mouse aortic endothelial cells were treated with glucose oxidase and flow cytometry was used to quantitate (A) ICAM-1, (B) I-Ab, (C) IsoLG, and (D) IsoLG-Adduct-H-2D^b^ FRET. Following infusion of sham or Ang II aortic ECs were evaluated for IsoLG-Adduct-H-2D^b^ FRET (E) Quantitation of FRET^+^ ECs. Data were analyzed by a two-tailed Student’s T-test or Mann-Whitney U Test (n = 6) (F) Representative histogram of FRET in aortic ECs. CD8^+^ T cells isolated from sham or AngII treated mice were co-cultured with GO or control treated mouse aortic endothelial cells and evaluated for proliferation. (G) Quantitation of proliferation of CD8^+^ T cells. Data were analyzed by a one-way ANOVA with Tukey’s post-hoc test (n = 6). (H) Representative histogram of CFSE staining of CD8^+^ T cells following co-culture.

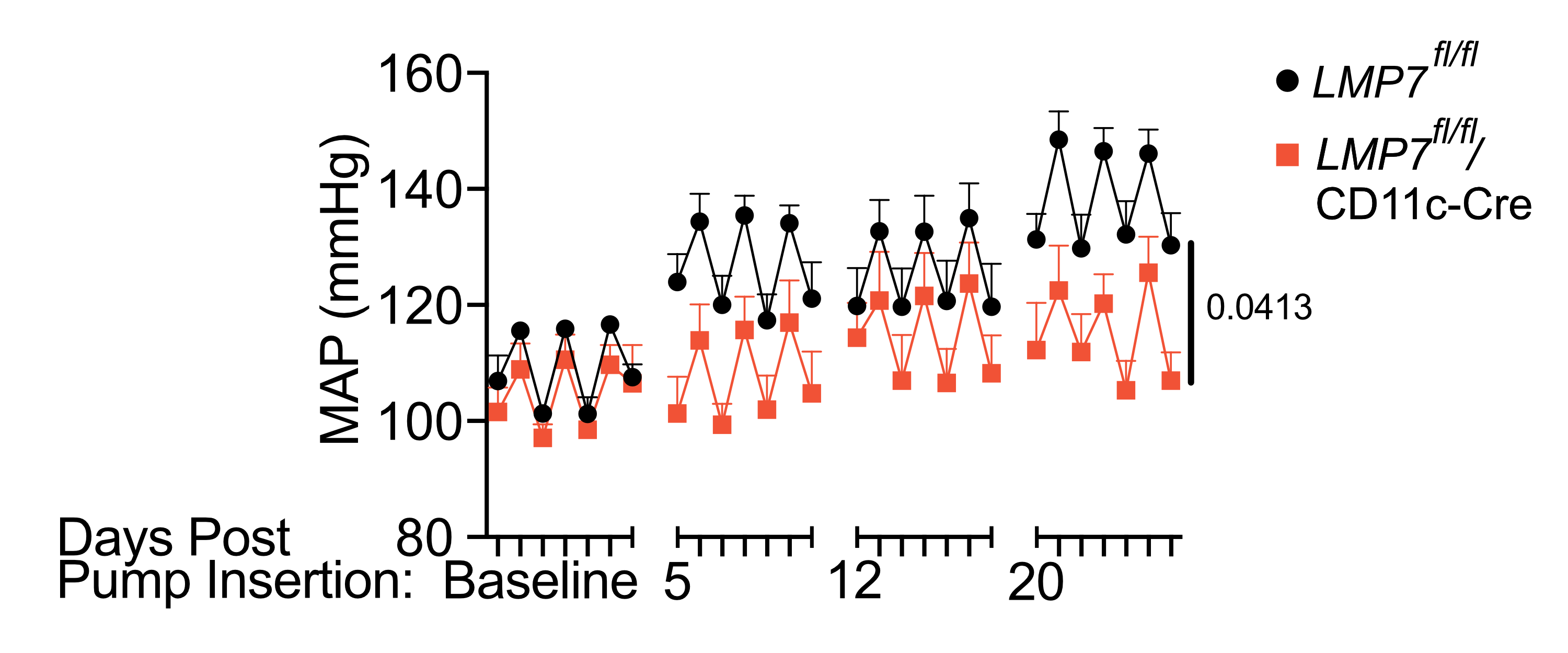

**Supplemental Figure 4. DC-specific *Lmp7* expression contributes to hypertension:** Radiotelemetry was used to measure mean artierial blood pressure. Blood pressure was analyzed using mixed-effects analysis (*n* = 5-6).

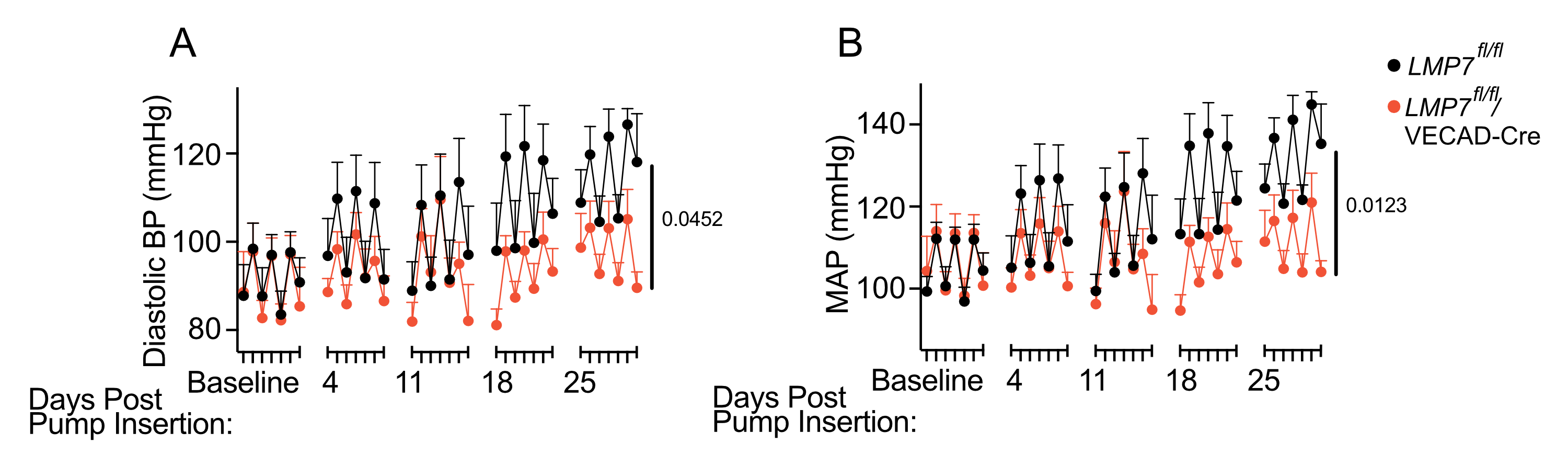

**Supplemental Figure 5. EC-specific *Lmp7* expression contributes to hypertension:** Radiotelemetry was used to measure (A) diastolic blood pressure and (B) mean arterial blood pressure (MAP). Blood pressure was analyzed using 2-way ANOVA (*n* = 5).

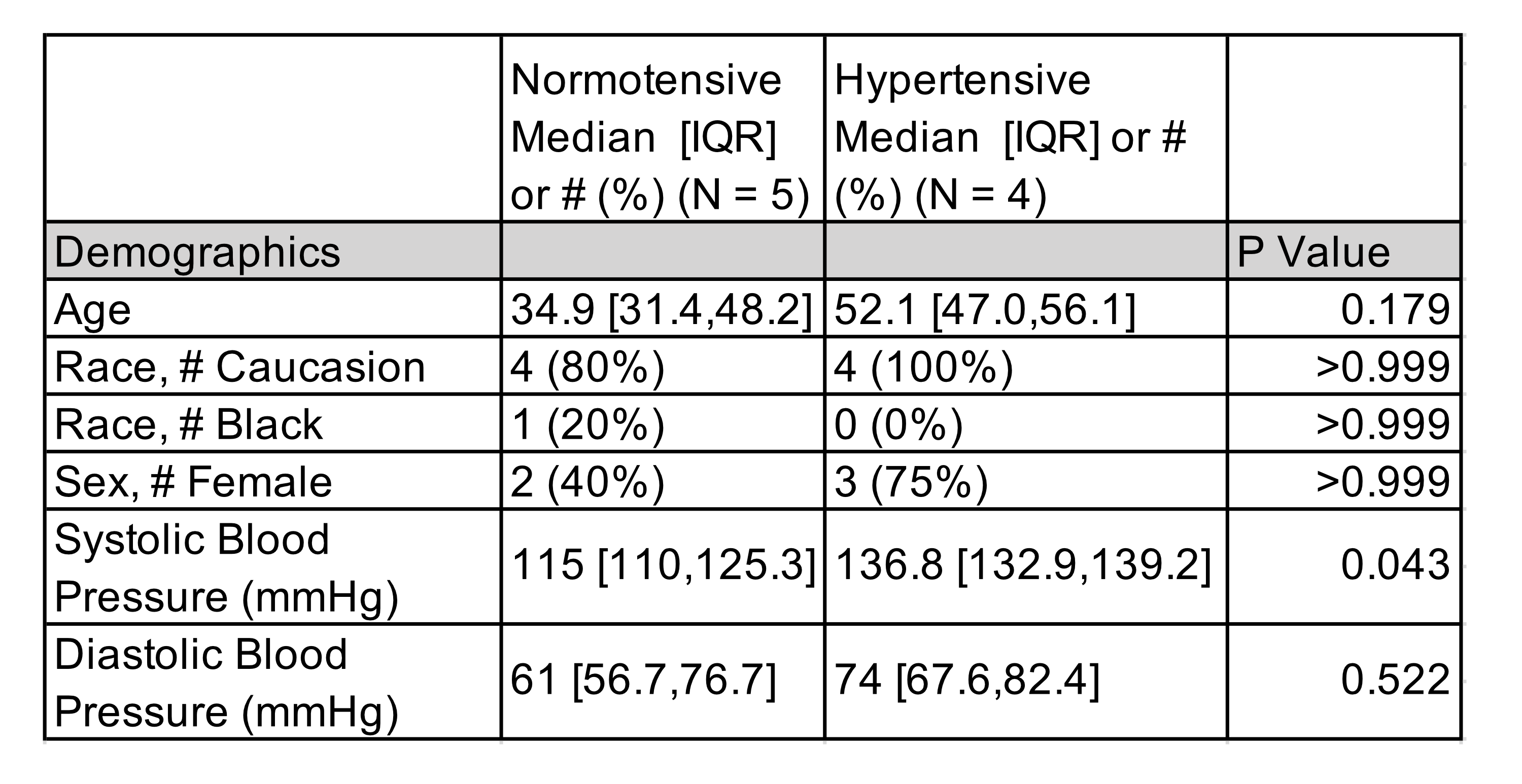

**Supplemental Table 1: Features of normotensive and hypertensive patients studied for LMP7 expression.** Data was compared with 2-way ANOVA with Bonferroni post-hoc test.
