## Supplementary material for "Immunoproteasomal Processing of Isolevuglandin Adducts in Hypertension": Key Resources Table

| **REAGENT or RESOURCE** | **SOURCE** | **IDENTIFIER** |
| --- | --- | --- |
| **Antibodies** | | |
| LMP7 | Abcam | Cat# ab3329;  RRID: AB_303708 |
| LMP7 | Cell Signaling | Cat# 13635S  RRID:AB_2744693 |
| β5 | Abcam | Cat# ab3330  RRID:AB_303709 |
| phospho-STINGser36 | Cell Signaling | Cat# 72971S  RRID:AB_2799831 |
| STING | Cell Signaling | Cat# 50494S  RRID:AB_2799375 |
| MHC-I H2 Db/D1 | Abcam | Cat# ab25244  RRID:AB_448732 |
| β-actin-Peroxidase | Sigma | Cat# A5441  RRID:AB_476744 |
| GAPDH | abcam | Cat# ab8245  RRID:AB_2107448 |
| CD31 AF594 | Biolegend | Cat# 102520  RRID:AB_2563319 |
| ICAM APC/Fire | Biolegend | Cat# 116125  RRID:AB_2716073 |
| IA/IE PerCP/Cy5.5 | Biolegend | Cat# 107625  RRID:AB_2191072 |
| D11-FcFv (anti-IsoLG-Lysine adduct) | GeneScript Biotech | N/A |
| CD45 APC | Invitrogen | Cat# 17-0451-82  RRID:AB_469392 |
| H-2Ld/H-2Db BV605 (MHC-I) | BD Biosciences | Cat# 742470  RRID:AB_2740804 |
| CD45 BV750 | Biolegend | Cat# 103157  RRID:AB_2734155 |
| CD3 BV605 | Biolegend | Cat# 100237  RRID:AB_2562039 |
| CD19 BV510 | Biolegend | Cat# 115545  RRID:AB_2562136 |
| CD4 FITC | BD Biosciences | Cat# 557307  RRID:AB_396633 |
| CD8 BV421 | Biolegend | Cat# 100738  RRID:AB_11204079 |
| MerTK APC | Biolegend | Cat# 151508  RRID:AB_2650739 |
| CD64 PE | Biolegend | Cat# 161003  RRID:AB_2904306 |
| CD11c BV650 | Biolegend | Cat# 117339  RRID:AB_2562414 |
| IA/IE PerCP/Cy5.5 | BD Biosciences | Cat# 562363  RRID:AB_11153297 |
| CD45 BB515 | BD Biosciences | Cat# 564590  RRID:AB_2738857 |
| MHC Class II (I-A/I-E) AF532 | ThermoFisher | Cat# 58-5321-82  RRID:AB_2811913 |
| CD11c AF594 | Biolegend | Cat# 117346  RRID:AB_2563323 |
| MerTK APC | Biolegend | Cat# 151508  RRID:AB_2650739 |
| CD3 PE/Cy7 | Biolegend | Cat# 100220  RRID:AB_1732057 |
| CD4 APC | BD Biosciences | Cat# 553051  RRID:AB_398528 |
| **Chemicals, peptides, and recombinant proteins** | | |
| Bortezomib (BTZ) | LC Laboratories | Cat# B-1408 |
| PR-957 | Apexbio | Cat# A4011 |
| MG-132 | Cell signaling | Cat# 2194S |
| Betadex Sulfobutyl Ether | USP | Cat# 1065550 |
| Sodium Citrate | Millipore-Sigma | Cat# PHR1416 |
| Glucose Oxidase | Millipore-Sigma | Cat# G7141-10KU |
| Ethyl-2-hydroxybenzlamine (Et2HOBA) | Vanderbilt University Medical Center | N/A |
| Angiotensin II (Ang II) | Millipore-Sigma | Cat# A2900 |
| Collagenase A | Millipore-Sigma | Cat# 10103586001 |
| Collagenase B | Millipore-Sigma | Cat# 11088815001 |
| Collagenase D | Millipore-Sigma | Cat# 10088858001 |
| LIVE/DEAD Fixable Violet Cell Stain | Thermo-Fisher | Cat# L34955 |
| Zombie NIR FIxabe Viability Kit | Biolegend | Cat# 423105 |
| DNAse I | Millipore-Sigma | Cat# 10104159001 |
| Luperox TBH70X, tert-Butyl hydroperoxide solution | Millipore-Sigma | Cat# 458139 |
| Streptavidin APC/Cy7 | Biolegend | Cat# 405208 |
| **Critical commercial assays** |  |  |
| ProteasomeGlo Assay | Promega | Cat# G8531 |
| CD11c MicroBeads UltraPure, mouse | Miltenyi | Cat# 130-125-835 |
| CD14 MicroBeads, human | Miltenyi | Cat#130-050-201 |
| Pan T cell Isolation Kit II, Mouse | Miltenyi | Cat#130-095-130 |
| Biotinylation Kit | Abcam | Cat# ab201795 |
| APEX Alexa Fluor 647 Antibody Labeling Kit | ThermoFisher | Cat# A10475 |
| CellTrace CVSE Cell Proliferation Kit | ThermoFisher | Cat# C34554 |
| **Deposited data** |  |  |
| RNA Sequencing Data | Geo Browser | GSE228902 |
| **Experimental models: Cell lines** |  |  |
| MAECs (Mouse Aortic Endothelial Cells) | Cell Biologics | Cat#C57-6052 |
| B8 fibroblasts | Marcus Groettrup | N/A |
| B8.27M.2 fibroblasts | Marcus Groettrup | N/A |
| **Experimental models: Organisms/strains** |  |  |
| Immunoproteasome Triple Knockout Mouse (TKO) | Regeneron Therapeutics |  |
| Mouse: *Lmp7^fl/fl^* mice | Gene Edit Biolab | N/A |
| Mouse: B6.Cg-Tg(Itgax-cre)1-1Reiz/J (CD11c-Cre) | Jackson Laboratory | Strain#008068 |
| Mouse: B6.FVB-Tg(Cdh5-cre)7Mlia/J (VE-Cadherin Cre) | Jackson Laboratory | Strain#006137 |
| Mouse: C57BL/6J | Jackson Laboratory | Strain#000664 |
| **Software and algorithms** |  |  |
| Gene Set Enrichment Analysis (GeSEA) | WebGestalt | https://www.webgestalt.org/ |
| FlowJo | BD Biosciences | N/A |
| Prism | Graphpad | N/A |
| ImageJ | NIH | N/A |
